# The molecular and cellular underpinnings of human brain lateralization

**DOI:** 10.1101/2025.04.11.648388

**Authors:** Loïc Labache, Sidhant Chopra, Xi-Han Zhang, Avram J. Holmes

**Author notes:** Corresponding authors: Loïc Labache.

## Abstract

Hemispheric specialization is a fundamental characteristic of human brain organization, where most individuals exhibit left-hemisphere dominance for language and right-hemisphere dominance for visuospatial attention. While some lateralized functions are evident in other species, the human brain displays a strong, species-wide bias. Despite the evolutionary and functional significance of these asymmetries, their molecular and cellular foundations remain poorly understood. Here, we identify key neurochemical and cellular asymmetries that underpin cortical lateralization. Specifically, we demonstrate lateralized gradients in neurotransmitter receptor densities, particularly along the multimodal monoaminergic-cholinergic axis, as well as asymmetries in mitochondrial distribution and the spatial prevalence of microglia and glutamatergic excitatory neurons. Using a multimodal approach that integrates in vivo functional MRI, PET imaging, and post-mortem transcriptomic and cellular data, we delineate two distinct cortical clusters: a left-lateralized network centered on language processing and a right-lateralized network supporting visuospatial attention. These results highlight a biologically embedded substrate for lateralized cognition that may inform both evolutionary theory and our mechanistic understanding of neuropsychiatric illnesses characterized by disrupted lateralization.

---

**H**emispheric specialization is a fundamental principle of human brain organization (1). Although lateralized functions have been observed in non-human animals (2, 3), the vast majority of humans exhibit left-hemispheric dominance for language and motor control (4, 5) and right-hemispheric dominance for visuospatial processing and spatial attention (6, 7). This complementary pattern likely reflects adaptive evolutionary pressures (8, 9), partly supported by interhemispheric connections via the corpus callosum (10–12). Although cerebral lateralization is theorized to be critical for the refinement of language and cognition across our evolutionary lineage, the precise origins and mechanisms underlying these pronounced functional asymmetries continue to be the subject of intense debate (13–18). Across development, the maturation of lateralized functions enhances visuospatial and language abilities while improving cognitive efficiency (19), with early anatomical markers of hemispheric specialization appearing during gestation (20) followed by a shift from intra- to inter-hemispheric connectivity at birth (20). This developmental trajectory continues through adolescence, as the transition from childhood to adulthood involves a reorganization of the attentional network’s connectivity (21). As highlighted in a recent review by Ocklenburg and colleagues (22), deviations from typical patterns of functional lateralization are commonly observed in individuals with neurodevelopmental (23, 24), psychiatric (25), and neurological disorders, such as autism (26) and schizophrenia (27). Recent multimodal neuroimaging syntheses further emphasize that hemispheric specialization for language is shaped by task demands, methodological choices, and interindividual factors such as handedness, motivating the need for integrative frameworks that can bridge functional, neurochemical, and structural markers of lateralization (28). In this regard, the extent to which the lateralization of discrete cognitive processes may be reflected in the molecular and cellular organization of the brain remains an open question with importance for the study of human behavior across both health and disease.

Regional variation in neurotransmitter receptor densities shape the functional specialization of the brain (29), broadly following the topography of large-scale networks amenable for study through functional magnetic resonance imaging (fMRI). This multi-scale organizational motif is evident in spatial associations between receptor densities and the hierarchical flow of sensory information (30) across cortex (31), suggesting that cerebral chemoarchitecture may reflect general organizational properties of brain structure and function (32). However, while converging evidence suggests that macro-scale functional systems are spatially coupled to specific receptor profiles, little is known about the neurotransmitter basis of task-related functional networks lateralization, such as those governing language and visuospatial attention. Despite the brain’s general bilateral symmetry (33), portions of the language network exhibit a neurotransmitter profile distinct from other cortical territories (34): certain left-hemisphere language regions involved in sentence comprehension exhibit elevated densities of serotonergic (5-HT_1A_) and cholinergic (M_2_) receptors, pointing to a potential neurotransmitter-based basis. Clinical and non-human animal work indicates that the left hemisphere may be shaped, at least in part, by a dopamine-driven system, supporting complex motor and language processing, while the right hemisphere seems to be modulated by a noradrenergic arousal system associated with visuospatial attention (35–37). These findings suggest that neurotransmitter distribution may not be symmetrical, and that a structured neurochemical axis could underlie the functional asymmetry observed in human cognition.

At the structural level, cortical thickness asymmetries have been linked to neurotransmitter organization (38), while lateralization in the distribution of neurotransmitters, such as GABA, has been observed (39). Of note, higher synaptic density in specific cortical regions correlates with improved cognitive performance (40), suggesting that the intricate organization and density of synaptic networks underlie not only basic sensory and motor functions but also higher-order cognitive processes. The functional properties of the human brain are reflected in the relative densities of cell types that pattern the cerebral cortex. For example, the functional organization of cortex as determined with intrinsic (resting-state) fMRI aligns with the spatial variability in its cellular composition as measured in post-mortem tissue (41–43). Indicating that these broad relationships may extend into properties of cortical lateralization, region-specific patterns of laminar gene expression within the left-hemisphere language network further underpin its structural and functional specialization (44). Another key factor that likely underpins the development of regional specialization is mitochondrial function, which is essential for meeting the brain’s high energy demands (45). Brain regions that serve as connectivity hubs exhibit enhanced mitochondrial respiratory capacities (46), supporting efficient cognitive processing, while even subtle disruptions in glucose supply can impair neural efficiency (47). Recent evidence links the intrinsic functional organization of the cortex to the spatial alignment of energy consumption, where regions with stronger functional connectivity exhibit greater metabolic demand (48). This reinforces the role of energy management in cognitive function. However, the extent to which spatial variability in cellular and mitochondrial functions mirrors the core properties of hemispheric lateralization remains unclear.

Here, using a diverse set of multimodal in vivo and post-mortem brain maps (29, 40, 41, 46, 49, 50) we investigate the spatial convergences that link functional, cellular, and molecular lateralization, and how these patterns might constrain the neural architecture underlying cognition. First, by combining three complementary language tasks (49) (production, reading, and listening), one visuospatial attention task (50), assessed through fMRI, and 19 neurotransmitter asymmetries maps (29), we establish that a multimodal monoaminergic-cholinergic axis underlies functional hemispheric specialization. Second, we provide evidence that mitochondrial phenotypes, alongside a specific population of intratelencephalic-projecting neurons and macrophages, are spatially coupled to this multimodal monoaminergic-cholinergic axis. Finally, we reveal that the brain is intrinsically organized around two opposing systems at the molecular and cellular levels: a left-lateralized system specialized for language processing and a right-lateralized system supporting visuospatial attention. In doing so, our analyses establish a direct link between microstructural brain organization and its macroscopic functional architecture, highlighting a tight relationship between in vivo measures of hemispheric functional specialization and its cellular and molecular substrates.

## Results

### A Multimodal Monoaminergic-Cholinergic Axis Underlines Functional Lateralization

To assess functional lateralization, we used three complementary language tasks (production, reading, and listening), and one visuospatial attention task. For these four tasks, we used the average fMRI activation map of 125 individuals with strong typical left-hemisphere language dominance, as defined by Labache and colleagues (4) from the BIL&GIN database (51), to reduce inter-individual variability. The 19 normative neurotransmitter density maps were derived from a databank compiling PET acquisitions from over 1,200 healthy individuals across nine neurotransmitter systems (29). Both the group-level fMRI and PET maps were parcellated into 326 cortical regions using a homotopic functional brain atlas optimized to study functional brain asymmetries (AICHA atlas (52)). Both fMRI and PET asymmetry maps were then computed by subtracting the regions in the right hemisphere from those in the left (Fig. 1, see Brain Maps – Functional Homotopic Atlas in the Methods).

**Figure 1.**
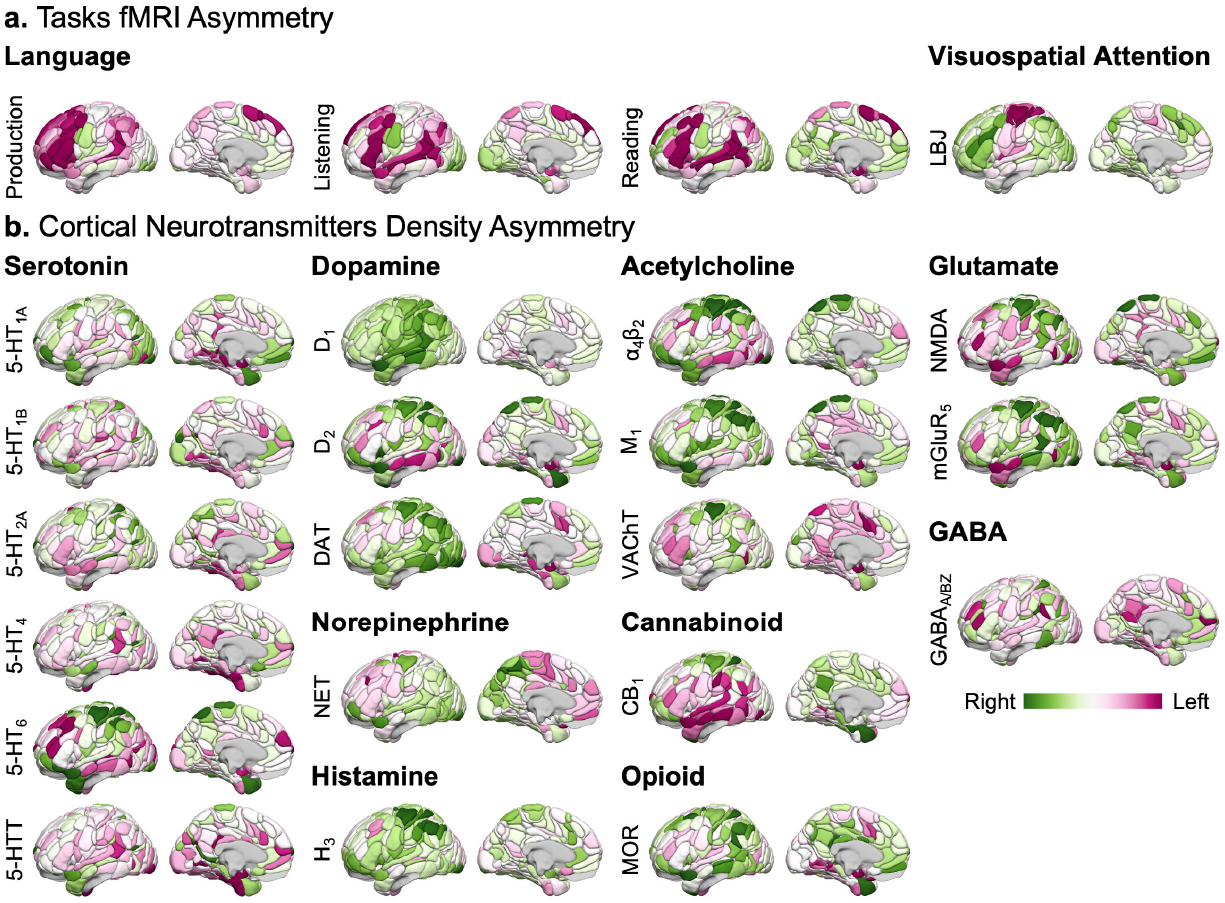
Asymmetry of normalized neurotransmitter densities and task-based fMRI. **a**, Set of four fMRI asymmetry maps defining supramodal language areas (49) (production, listening, reading, contrast: sentence minus list of words) and visuospatial attention (50) (LBJ: Line Bisection Judgment). The maps were collated and averaged across 125 healthy participants from the *BIL&GIN* database (51), previously identified as typically brain-organized for language (4) (*i*.*e*. left hemisphere dominant). Together, these four tasks assess the hemispheric complementarity of language and attention. **b**, 19 PET asymmetry maps of neurotransmitter receptor and transporter density collated by Hansen and colleagues (29). This set covers nine neurotransmitter systems and includes data from 1,238 participants. The maps encompass 163 cortical regions defined by the homotopic AICHA atlas (52). Asymmetries were calculated by subtracting left hemisphere values from right hemisphere values and are visualized on the left hemisphere. The 3D rendering in the MNI space was created using Surf Ice software (https://www.nitrc.org/projects/surfice/).

The 19 neurotransmitter maps captured receptors and transporters from nine neuromodulatory systems (Fig. 1b), each exhibiting characteristic hemispheric asymmetries. Glutamatergic markers showed system-specific patterns, with NMDA receptors exhibiting mixed left- and right-lateralized clusters across frontal, temporal, and parieto-occipital regions, while mGluR_5_ was largely right-lateralized except in parahippocampal and superior temporal areas. GABAergic density was predominantly left-lateralized, most notably in anterior cingulate and lingual cortices. Opioid and cannabinoid systems showed heterogeneous topographies, with distinct left- and right-lateralized clusters distributed across temporal, frontal, and parietal cortices. Cholinergic receptors displayed shared patterns of left-lateralization in temporal and midcingulate regions and right-lateralization elsewhere, whereas VAChT showed a broader leftward shift. Histamine was globally right-lateralized, and norepinephrine showed left-lateralized frontal-paracentral regions contrasted with more right-lateralized posterior cortices. Dopaminergic markers exhibited complementary asymmetries across lateral versus medial surfaces, while serotonergic receptors showed consistent but subtype-specific combinations of leftward and rightward asymmetry across temporal, frontal, and occipital regions. Together, these normative maps provide a compact, multireceptor characterization of neurochemical lateralization across the cortex.

To investigate the link between cortical lateralization with neurotransmitter density asymmetries, we used permutation-based Canonical Correlation Analysis (CCA) (53). This method identifies linear combinations of variables from two distinct feature sets (in this case, cortical lateralization and neurotransmitter density asymmetries) that maximize the correlation between them. Using CCA, we discovered pairs of canonical variables (modes) that reveal the extent to which multivariate neurotransmitter profiles and functional lateralization patterns for language and visuospatial attention co-vary across corresponding homotopic regions within each hemisphere. Before conducting CCA, we used Principal Component Analysis (PCA; Extended Data Fig. 1) to produce a parsimonious and interpretable description of the original datasets (54, 55). To account for spatial autocorrelation between brain maps when testing for statistical significance (56), we used “spin tests,” where cortical regions are repeatedly rotated on an inflated sphere to create null configurations that maintain the cortex’s spatial autocorrelation. These null maps were then used to generate a null distribution of canonical correlation coefficients (10,000 permutations), allowing us to assess the statistical significance of the observed CCA modes. Next, we identified the specific functional and neurotransmitter features driving each CCA mode by computing canonical loadings, correlating each input map with its corresponding canonical variate, and assessing their reliability through bootstrap-derived z-scores and Holm-corrected significance testing.

The CCA revealed a significant set of modes underlying the relationship between functional lateralization and neurotransmitter density asymmetry. These showed a significant relationship between functional cortical lateralization and neurotransmitter density asymmetry (*r*=0.39, *p*_spin_=0.0058, Fig. 2). The neurotransmitter density asymmetry mode, hereafter referred to as the neurotransmitter asymmetry axis, explained 21% of the variance across the 19 neurotransmitter variables (Fig. 2a). The task fMRI asymmetry mode, hereafter referred to as the task asymmetry axis, explained 24% of the variance of the four functional variables (Fig. 2b). To identify the neurotransmitters and tasks that contribute most to each mode, we computed loadings by correlating each input variable with the corresponding canonical mode (Fig. 2a-b). The resulting Pearson loadings (Extended Data Table 1) indicated that the neurotransmitter asymmetry axis was anchored on its positive end by acetylcholine (M_1_) receptors (*r*=0.80, *p*_FWER_=9.10^−7^), and on its negative end by norepinephrine (NET) receptors (*r*=−0.18, *p*_FWER_=1). However, this mode also reflects substantial contributions from other systems, including dopaminergic (e.g., D_1_: *r*=0.71, *p*_FWER_=8.10^−11^, DAT: *r*=0.61, *p*_FWER_=2.10^−5^, D_2_: *r*=0.60, *p*_FWER_=6.10^−3^), and histaminergic (H_3_: *r*=0.70, *p*_FWER_=1.10^−4^), as well as more acetylcholinergic receptors (α_4_β_2_: *r*=0.55, *p*_FWER_=3.10^−2^, VAChT: *r*=0.47, *p*_FWER_=1.10^−3^), underscoring its multivariate neurochemical nature. All the lateralized task variables were negatively and significantly correlated with the task fMRI asymmetry canonical mode, with the listening task showing the strongest negative correlation (*r*=−0.60, *p*_FWER_=6.10^−3^), followed by the reading task (*r*=−0.52, *p*_FWER_=0.028), production (*r*=−0.50, *p*_FWER_=0.046), and attention (*r*=−0.49, *p*_FWER_=0.048). Meaning that during language-related tasks, the classically observed leftward-lateralized regions co-vary with a right-lateralized density of the muscarinic acetylcholine receptor M_1_ and a left-lateralized density of the norepinephrine NET receptor. Conversely, we observed the opposite pattern in right-lateralized visuospatial attention tasks, where these receptor densities were reversed. Given its composition; a linear combination of 19 neurotransmitter maps with significant contributions from cholinergic and multiple monoaminergic systems, we refer to this multivariate pattern as the multimodal monoaminergic-cholinergic axis, reflecting the dominant influence of M_1_ acetylcholine receptors together with significant dopaminergic and histaminergic components. The anchor of this axis aligns with prior work showing that acetylcholine interacts with dopamine (top five contributors to the neurotransmitter mode, see D_1_, DAT, and D_2_ in Fig. 2a) to support complex motor programming, especially articulatory speech motor control, while norepinephrine modulates arousal and attention (35). However, neurotransmitter laterality relationships are inherently complex and reflect multimodal interactions across a diverse set of receptor and transmitter types. Clinically, norepinephrine modulating drugs may facilitate recovery from aphasia by enhancing synaptic plasticity (57). Similarly, acetylcholine-targeting drugs may offer comparable therapeutic benefits (57). These observations suggest the importance of maintaining a balance between acetylcholine and norepinephrine receptor activity. The second set of modes was not significant (*r*=0.17, *p*_spin_=0.3274, Extended Data Fig. 2c). A full description of the second set of modes can be found in Extended Data Fig. 3.

**Figure 2.**
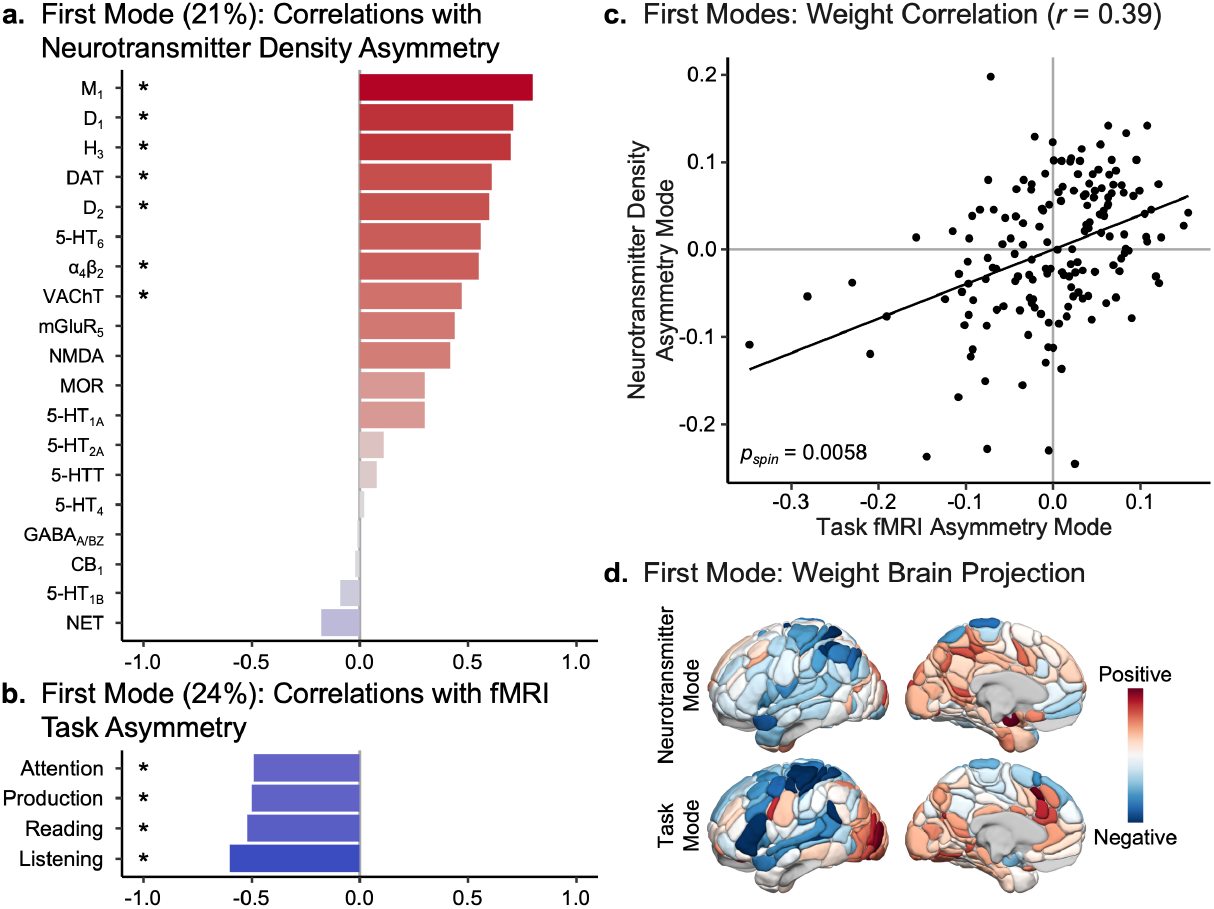
Canonical Correlation Analysis (CCA) results reveal the neurotransmitter receptor and transporter substrates underlying functional cortical lateralization. **a**, Correlation values of the neurotransmitter mode with the 19 normalized neurotransmitter density asymmetries. This neurotransmitter mode explained 21% of the variance in the normalized neurotransmitter density asymmetry maps. An acetylcholine (muscarinic acetylcholine receptor M_1_, M_1_ receptor) versus norepinephrine (norepinephrine transporter, NET receptor) axis is identified, indicating that a positive value of the neurotransmitter mode corresponds to a leftward asymmetry in acetylcholine M_1_ receptor density and a rightward asymmetry in norepinephrine NET receptor density. Asterisks (*) indicated statistical significance using bootstrapping (*p*_FWER_<0.05). **b**, Correlation values of the task mode with the four functional asymmetry maps. This task mode explained 24% of the variance in task asymmetry. A positive value of the task mode corresponds to a leftward asymmetry during the attention, listening, reading, and production tasks. Asterisks indicated statistical significance using bootstrapping (*p*_FWER_<0.05). **c**, Scatter plot showing the association between neurotransmitter and task modes. The neurotransmitter and task modes are significantly correlated (*r*=0.39, *p*_spin_=0.0058; significance was assessed using a spin test with 10,000 permutations (60). This significant covariation indicates that the leftward asymmetry displayed in the supramodal language network (49) during language tasks is associated with leftward asymmetry in acetylcholine receptor density and rightward asymmetry in norepinephrine receptor density. An opposite receptor relationship underlies the rightward asymmetries observed in the temporo-frontal and parieto-frontal visuospatial attentional networks (50). **d**, Projection of each regional CCA weight on the 163 cortical regions defined by the homotopic AICHA atlas (52). In the top rows, the positive red regions are associated with leftward asymmetry of acetylcholine receptor density, while the negative blue regions correspond to rightward asymmetry of norepinephrine receptor density. In the bottom rows, the positive red regions are associated with rightward lateralization during the four tasks, whereas the negative blue regions are associated with leftward asymmetries.

To identify the brain regions contributing most significantly to the relationship between neurotransmitter asymmetry and task asymmetry, we projected the canonical weights from both the neurotransmitter asymmetry axis and the task asymmetry axis back onto the brain (Fig. 2d) and examined the top 10% of regional weights. For the task mode, the regions with negative weights, corresponding to leftward asymmetries, overlapped at 41% with the core language network (49), 28% with the attention networks (50), and 26% with somato/motor areas supporting manual preference (50, 58). Notably, these regions included the frontal gyrus (*pars triangularis*) and the superior temporal sulcus; both defined as resting-state hubs for the language network (49), as well as the postcentral sulcus and parts of the Rolandic fissure, which are somato/motor hubs at rest (50), and the inferior frontal sulcus, a key region in the parieto-frontal attention network (50). Positive weights, indicating rightward lateralization, corresponded to 24% of regions supporting visuospatial attention processing (50), and 29% to the visual network (50, 59). Regarding the neurotransmitter asymmetry axis or multimodal monoaminergic-cholinergic axis, 12% of the most negative weights overlapped with the language network, 18% with the attention network, and 29% with the somato/motor network, including a hub (postcentral sulcus). This is consistent with prior work suggesting that M1 receptors are higher in the left inferior frontal gyrus and superior temporal gyrus (34), regions overlapping with the language (49), and attention networks (50) and showing the highest concentration in M1 receptors (31). Conversely, positive weights were associated with the visual network (12%), and the somato/motor network (6%). These findings highlight a clear relationship between asymmetries in neurotransmitter densities and the functional specialization of brain networks, suggesting that the spatial heterogeneity in neurotransmitter distribution may contribute to the diversity of functional specializations across the cortical sheet (34).

### Mitochondrial Phenotypes as Substrates for the Lateralization of the Multimodal Monoaminergic-Cholinergic Axis

At the molecular level, mitochondria generate cellular energy through ATP production and regulate key cellular processes, allowing adaptation to metabolic demands (61). They are critical in energy-intensive tissues such as the brain, where they balance neuronal excitability and neurotransmitter release, modulate synaptic plasticity (45), and exhibit spatial expression patterns that closely align with the evolutionary development of the brain (46). Moreover, highly connected regions demand higher energy levels (62, 63) and play crucial roles in cognition (64, 65). The density of synapses, fundamental to information processing, have recently been shown to reflect functional topography and mirror hierarchical functional network organization (40). Furthermore, brain regions with higher synaptic density exhibit greater mitochondrial metabolism suggesting mitochondria critically support synaptic activity and integrity across the human brain (66, 67). However, a direct link between these molecular properties and the lateralization of fundamental brain functions, such as language and attention, has yet to be established.

Here, we investigated how synaptic and mitochondrial activities underpin the multimodal monoaminergic-cholinergic axis, supporting language lateralization and visuospatial attention. Until recently, whole-brain maps of synaptic density and mitochondrial function were not achievable. Advances in PET imaging of synaptic vesicle glycoprotein 2A (SV2A) (40), an index of synaptic density, along with newly developed quantitative mitochondrial mapping methods (46); including tissue respiratory capacity (TRC), mitochondrial tissue density (MitoD), and mitochondrial respiratory capacity (MRC), now allow comprehensive analyses of these critical molecular systems. Linear regression was employed, using the multimodal monoaminergic-cholinergic axis as the dependent variable. Significance was assessed using a spin test (56) with 10,000 null rotations to control for spatial autocorrelation. P-values were corrected for multiple comparisons using the false discovery rate (FDR) method (68), accounting for the four regression models applied separately for each molecular map (SV2A, TRC, MitoD, and MRC) assessing their contribution to variation along the multimodal monoaminergic-cholinergic axis. As with the neurotransmitter maps, the molecular maps were parcellated into 326 cortical regions from the AICHA atlas, and asymmetry maps were computed by subtracting the right regions from the left ones (Fig. 3a).

**Figure 3.**
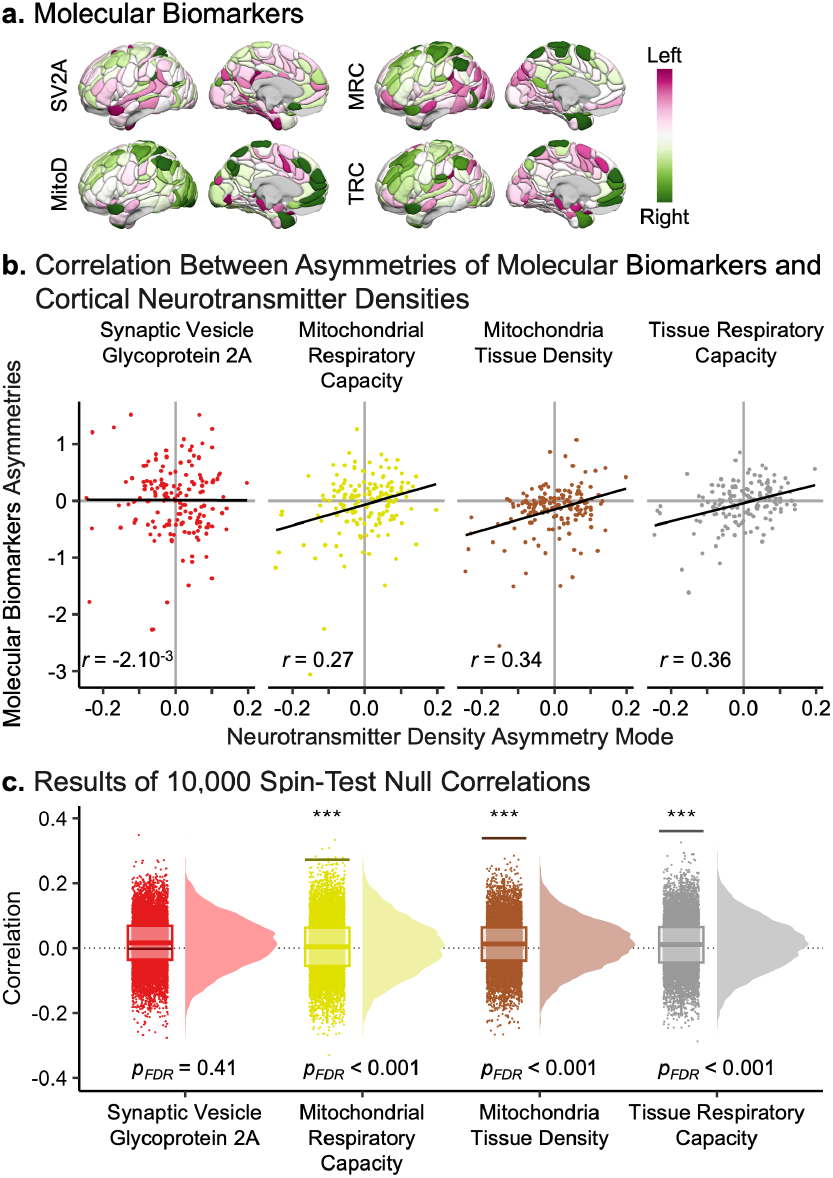
Molecular substrates of the neurotransmitter receptor and transporter mode underlying functional cortical lateralization. **a**, Set of four normalized asymmetry maps of molecular biomarkers. SV2A represents the asymmetry of synaptic density (Synaptic Vesicle Glycoprotein 2A), as collected by Johansen and colleagues from 33 healthy participants (40). TRC (Tissue Respiratory Capacity), MitoD (Mitochondria Tissue Density), and MRC (Mitochondrial Respiratory Capacity) correspond to three maps of mitochondrial activity collected by Mosharov and colleagues from a 54-year-old donor (46). Asymmetries were calculated by subtracting left hemisphere values from right hemisphere values (Right - Left) and are visualized on the left hemisphere. The maps encompass 163 cortical regions defined by the homotopic AICHA atlas (52). **b**, Correlation between the neurotransmitter mode and the asymmetries of the four molecular biomarkers. The neurotransmitter mode shows a significant association (after false discovery rate correction (68)) with all mitochondrial activity maps: TRC (*r*=0.36, *p*_FDR_<1.10^−4^), MitoD (*r*=0.34, *p*_FDR_<1.10^−4^), and MRC (*r*=0.27, *p*_FDR_=0.0009). The correlation between the neurotransmitter mode and synaptic density (SV2A) is not significant (*r*=0.00, *p*_FDR_=0.4069). **c**, Results of the spin tests (60) assessing the significance of the Pearson correlations between the neurotransmitter mode and the molecular biomarkers. The spin tests were performed using 10,000 permutations.

Linear regression analyses identified significant associations between mitochondrial phenotypes and the multimodal monoaminergic-cholinergic axis (Fig. 3b–c). Specifically, MRC was correlated (*r*=0.27, *p*_FDR_=0.0009), as were MitoD (*r*=0.34, *p*_FDR_<1.10^−4^) and TRC (*r*=0.36, *p*_FDR_<1.10^−4^). In contrast, the association between the multimodal monoaminergic-cholinergic axis and SV2A did not reach statistical significance (*r*=−2.10^−3^, *p*_FDR_=0.41; Fig. 3b–c). These findings suggest that a leftward asymmetry in mitochondrial phenotypes, but not synaptic density, corresponds with a leftward asymmetry in acetylcholine and a rightward asymmetry in norepinephrine neurotransmission. The strongest rightward asymmetry across the three phenotypes was located in the paracentral lobule. In contrast, the strongest leftward asymmetries differed among phenotypes: the superior temporal pole gyrus for TRC, the lingual gyrus for MitoD, and the supramarginal gyrus for MRC.

### Cellular Correlates of the Multimodal Monoaminergic-Cholinergic Axis Lateralization

While the previous analyses identified molecular-scale correlates of laterality, microscale features, such as specific cell types, may also constrain functional lateralization. Regional cellular profiles spatially covary with the macroscale functional organization of the cortical sheet, as estimated through fMRI (41). Moreover, language lateralization is reflected across the entire macroscale functional organization of the cortex (5). However, the direct link between these regional cellular fingerprints and language lateralization, and more broadly, cognitive functions such as language and attention, remains to be established.

We used a bidirectional stepwise linear regression based on the Akaike Information Criterion to examine the relationship between the asymmetry of these cellular fingerprints and the lateralization of language and attention. This analysis linked 24 lateralization maps of transcriptomically-defined cell-type densities and the neurotransmitter mode (multimodal monoaminergic-cholinergic axis) underlying the functional lateralization of language and attention. To control for spatial autocorrelation between cell-type maps and the neurotransmitter mode map, we utilized a spin test (56), generating null configurations by rotating cortical regions on an inflated sphere. These configurations (10,000) were used to shuffle the cellular fingerprints, allowing us to assess the individual statistical significance of each selected cell-type. The p-values were corrected for multiple comparisons using the FDR method (68).

As described by Zhang and colleagues (41), these 24 cellular classes exhibit distinct laminar specializations, developmental origins, morphologies, spiking patterns, and broad projection targets (69). The cell-types (Extended Data Table 2) include nine GABAergic inhibitory interneurons (PAX6, SNCG, VIP, LAMP5, LHX6, Chandelier, PVALB, SST CHODL, SST), nine glutamatergic excitatory neurons (L2/3 IT, L4 IT, L5 IT, L6 IT, L5 ET, L5/6 NP, L6 CT, L6b, L6 IT Car3), and six non-neuronal cells (Astro, Endo, VLMC, Oligo, OPC, Micro/PVM).

Following Zhang and colleagues, each cell-type’s abundance was estimated in available bulk samples and aggregated into the 163 cortical brain regions of the AICHA atlas. Asymmetries for each cell-type were computed by subtracting the standardized value of the right hemisphere from that of the left. Due to the ex vivo sampling technique, cell fingerprint asymmetries were available for only 123 brain regions. A complete map of the 24 standardized cell-types’ asymmetry is presented in Extended Data Fig. 4.

The bidirectional stepwise linear regression identified five cell-types: L5 IT (*β*=−0.018, *t*_*β*_=−3.27), Micro PVM (*β*=−0.014, *t*_*β*_=−2.36), L6 IT (*β*=−0.012, *t*_*β*_=−2.30), L2/3 IT (*β*=−0.014, *t*_*β*_=−2.17), and L6 IT Car3 (*β*=−0.008, *t*_*β*_=−1.39), out of the 24 examined (Fig. 4a-b) that significantly explain variance in the multimodal monoaminergic-cholinergic axis (*F*_model_=3.493, *p*_model_=0.0056, *R*^2^= 12.99%). Four of these significant cell-types remained associated with the multimodal monoaminergic-cholinergic axis balance after performing spin tests (Fig. 4c): L5 IT (*p*_FDR_=0.0030), Micro PVM (*p*_FDR_=0.0218), L6 IT (*p*_FDR_=0.0218), and L2/3 IT (*p*_FDR_=0.0218), while L6 IT Car3 is not significant (*p*_FDR_=0.0864). All cell-types contributed negatively to the multimodal monoaminergic-cholinergic axis, indicating that a rightward asymmetry for the selected cell-types is associated with a leftward asymmetry in acetylcholine and rightward asymmetry in norepinephrine neurotransmitters. Specifically, L5 IT denotes layer 5 intratelencephalic-projecting neurons which have shorter basal dendrites (ends at layers 2 and 3) and simple apical dendrites compared with ITs in the upper layers; Micro/PVM refers to non-neuronal microglia and perivascular macrophages; L6 IT is for layer 6 intratelencephalic-projecting neurons and tends to connect with local neurons in layer 6 reciprocally; L2/3 IT stands for layer 2 and 3 intratelencephalic-projecting neurons and its dendrites extend within upper layers, while its the axons target layer 5, finally L6 IT Car3 signifies layer 6 intratelencephalic-projecting (Car3-like), characterized by widespread intracortical axonal projections but an absence of collateral projections to the striatum.

**Figure 4.**
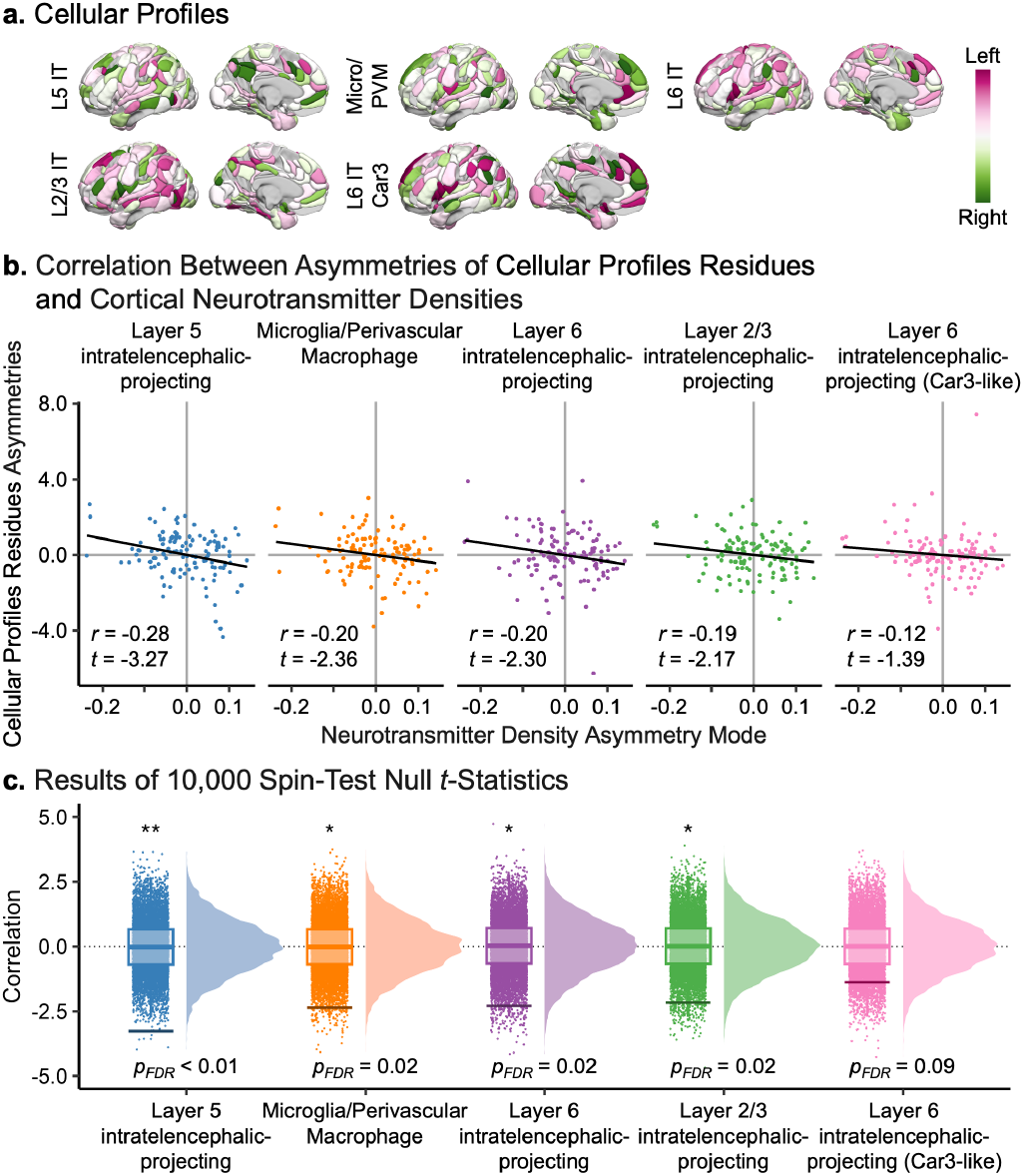
Cellular substrates of the neurotransmitter receptor and transporter mode underlying functional cortical lateralization. **a**, Set of five normalized asymmetry maps of cellular biomarkers as reported by Zhang and colleagues (41), derived from bulk samples of six postmortem brains (80). Asymmetries were calculated by subtracting right hemisphere values from left hemisphere values (left - right) and are visualized on the left hemisphere. The maps encompass 123 cortical regions defined by the homotopic AICHA atlas (52). These four cell-types were selected from 24 through bidirectional stepwise linear regression (based on the Akaike Information Criterion). The resulting linear model is significant (*F*_model_=3.493, *p*_model_=0.0056) and explains 12.99% of the variance of the neurotransmitter mode. The 24 asymmetry maps can be visualized in Extended Data Fig. 4. Layer 5 (L5) intratelencephalic (IT)-projecting neurons have shorter basal dendrites (ends at layers 2 and 3) and simple apical dendrites compared with ITs in the upper layers, Micro/PVM refers to non-neuronal microglia and perivascular macrophages, L6 IT tends to connect with local neurons in layer 6 reciprocally, L2/3 IT have their dendrites extend within upper layers, while their axons target layer 5, finally L6 IT Car3 signifies layer 6 intratelencephalic-projecting (Car3-like). **b**, Visualization of the bidirectional stepwise linear regression results through the correlation between the residuals of the cellular asymmetry profiles and the neurotransmitter mode. The residuals were obtained by regressing, for each cell-type, the effects of the other three cell-types, allowing visualization of the one-to-one relationship linking each cellular asymmetry profile to the neurotransmitter mode. The neurotransmitter mode shows a significant association (after false discovery rate correction (68)) with the following cell-types: L5 IT (*β*=−0.018, *t*_*β*_=−3.27, *p*_spin_=0.0006, *p*_FDR_=0.0030), Micro PVM (*β*=−0.014, *t*_*β*_=−2.36, *p*_spin_=0.0101, *p*_FDR_=0.2175), L6 IT (*β*=−0.012, *t*_*β*_=−2.30, *p*_spin_=0.0133, *p*_FDR_=0.2175), and L2/3 IT (*β*=−0.014, *t*_*β*_=−2.17, *p*_spin_=0.0174, *p*_FDR_=0.2175). The correlation between the neurotransmitter mode and L6 IT Car3 (*β*=−0.008, *t*_*β*_=−1.39, *p*_spin_=0.0864, *p*_FDR_=0.0864). **c**, Results of the spin tests (60) assessing the significance of the beta coefficients (*β*) associating the neurotransmitter mode and the cellular biomarkers. The spin tests were performed using 10,000 permutations.

Consistent with recent findings, L6 IT neurons are associated with somato/motor and ventral attention networks, supporting “bottom-up” attention (41). In contrast, L5 IT neurons link to transmodal regions (70), including dorsal attention and frontoparietal networks, and interact with layer 2/3 neurons in “top-down” processes (41). Micro/PVM cells are enriched in limbic networks, suggesting that these cells could influence the unique functional properties of limbic regions, such as their role in emotion and memory processing (41). Language network lateralization spans multiple macroscale gradients and associative networks (5), and the enrichment of these cell-types within these gradients further underscores their pivotal role in language function and hemispheric lateralization. Consistent with this organization, recent work shows that a left-lateralized orthographic visual stream converges onto a basal inferotemporal language region, forming a reading-specific nexus within the broader language network (71). Laminar asymmetry analyses (72) indicate that corticothalamic layer 6 cells exhibit distinct asymmetry patterns, shifting from leftward asymmetry in superficial layers to rightward asymmetry in deeper layers. Layer 5 also shows laminar-specific asymmetry in sensorimotor and temporal output regions, while layer 3 pyramidal neurons drive leftward asymmetries in areas such as the inferior frontal cortex.

L5 IT neurons showed strong rightward asymmetry in the cingulate sulcus and the middle frontal gyrus related to visuospatial attention processing (50) while showing strong leftward asymmetry in areas related to language (49) and somato/motor processing (50) (supramarginal gyrus and precentral sulcus). L5 IT neurons play a critical role in cortical sensory processing and produce internal models fed back to L5 ET neurons and are engaged in the preparatory stages of tasks, actively participating in sensory sampling and decision-making processes (73).

Micro/PVM exhibited pronounced rightward lateralization in the middle and superior temporal gyrus and the superior frontal gyrus; areas involved in language (49) and visuospatial attention (50). In contrast, Micro/ PVM cells were predominantly leftward lateralized in areas associated with manual preference (50, 58), such as the Rolandic operculum and the precentral sulcus, and visuospatial attention, such as the supramarginal and the anterior cingulum gyrus. These specialized brain macrophages maintain vascular and neural integrity, modulate immune responses, and adapt to local environments to support homeostasis and circuit assembly (74, 75).

L6 IT neurons were principally rightward lateralized in the temporo-frontal network related to attention processing (50), while being leftward asymmetrical in the somato/motor network (50, 58). L6 IT neurons are involved in local circuit modulation, integrating deep-layer signals and associative processes and fine-tuning cortical processing and feedback circuits (76, 77).

Conversely, L2/3 IT neurons showed strong rightward lateralization in the somato/motor network (50, 58) while being leftward dominant in the language network, notably in the superior temporal sulcus, a hub region for the supramodal language network (49). Layer 2/3 intratelencephalic-projecting neurons facilitate intracortical communication by integrating and transmitting information both locally and through long-range cortical networks (76), a key feature of the widespread language network (78), with feedforward or feedback roles depending on their cortical location (79).

We have demonstrated that mitochondrial phenotypes and cellular markers (L5 IT, Micro PVM, L6 IT, and L2/3 IT) significantly underlie the multimodal monoaminergic-cholinergic axis responsible for lateralizing language and visuospatial attention. Next, we examine how these cellular and molecular asymmetries present across canonical neural networks.

### Opposing Lateralization Profiles in Language- and Attention-related Networks at the Molecular and Cellular Level

Finally, we investigated the specificity of molecular and cellular fingerprints among seven canonical large-scale functional networks (59) as defined by Yan and colleagues (81), focusing on three significant molecular markers: TRC (tissue respiratory capacity), MitoD (mitochondrial tissue density), and MRC (mitochondrial respiratory capacity), and four cellular substrates: L5 IT (layer 5 intratelencephalic-projecting neurons), Micro/ PVM (microglia/perivascular macrophage cells), L6 IT (layer 6 intratelencephalic-projecting neurons), and L2/3 IT (layer 2 and 3 intratelencephalic-projecting neurons), that underlie functional lateralization (Fig. 5a–b). These seven biomarkers revealed the molecular and cellular organization of the seven canonical functional networks. Each network was characterized by its normalized values for the seven biomarkers.

**Figure 5.**
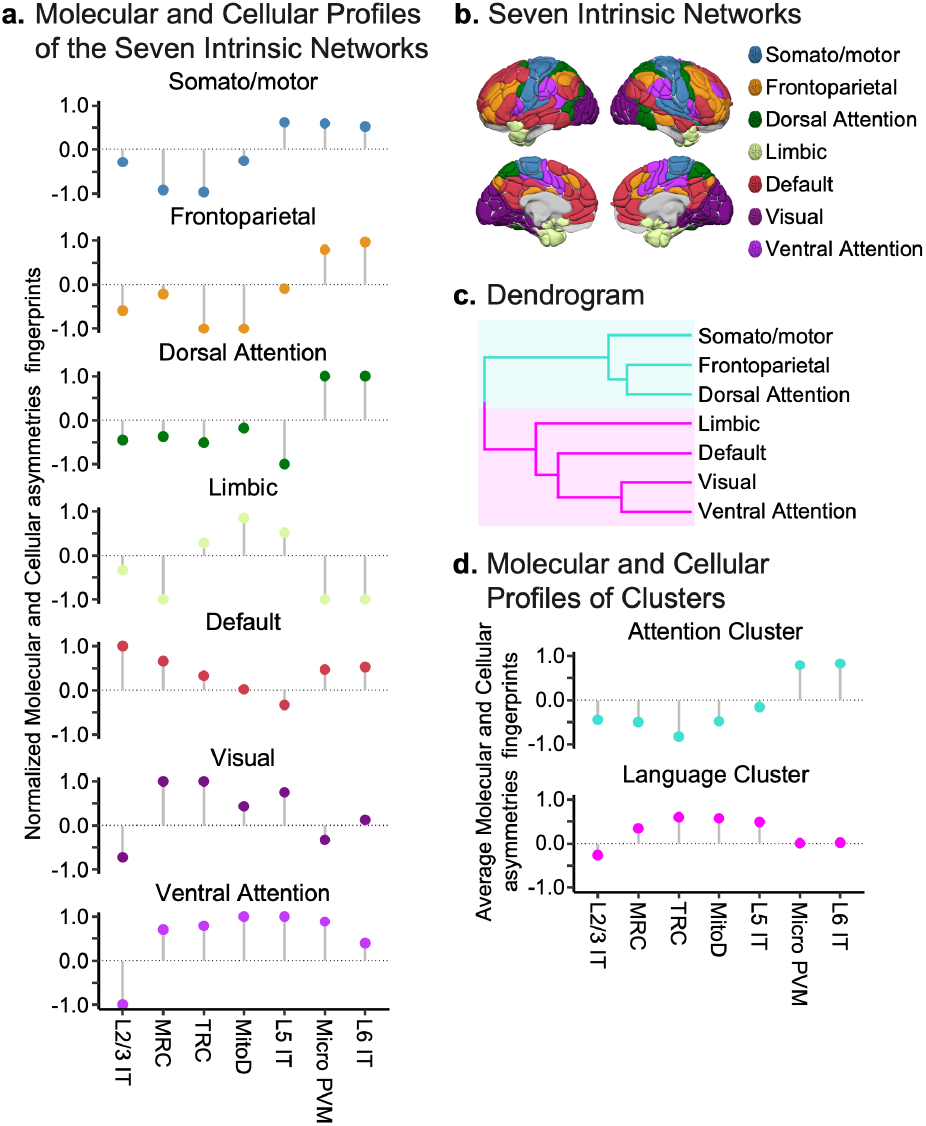
Organization of molecular and cellular profiles of the seven canonical intrinsic networks. **a**, Average significant normalized molecular (Mitochondrial Respiratory Capacity; MRC, Tissue Respiratory Capacity; TRC, Mitochondria Tissue Density; MitoD) and significant cellular (Layer 2 and 3 intratelencephalic-projecting neurons; L2/3 IT, layer 5 intratelencephalic-projecting neurons; L5 IT, Microglia/PeriVascular Macrophage cells; Micro/PVM, Layer 6 intratelencephalic-projecting neurons; L6 IT) profiles of the seven canonical intrinsic networks (59) as defined by Yan and colleagues (81). **b**, Brain organization according to the seven canonical intrinsic networks overlaid on the 246 parcels of the AICHA atlas (52). **c**, Hierarchical clustering identifies two clusters among the seven canonical intrinsic networks. The first cluster, shown in blue, is associated with attentional processes, as its regions overlap the task fMRI-defined lateralized visuospatial attention atlas (50) (*SDI*=0.33, Sørensen-Dice index). The second cluster, shown in pink, is associated with language processes, as its regions overlap the task fMRI-defined lateralized supramodal language atlas (49) (*SDI*=0.25). **d**, Molecular and cellular profiles of the clusters. The attentional cluster exhibits rightward lateralization for L2/3 IT, MRC, TRC, MitoD, L5 IT and leftward lateralization for Micro PVM and L6 IT. Conversely, the language cluster shows leftward lateralization for MRC, TRC, MitoD, L5 IT, Micro PVM and L6 IT and rightward lateralization for L2/3 IT.

Using the molecular and cellular fingerprints, we derived two clusters (Fig. 5c) via agglomerative hierarchical clustering (82), employing Euclidean distance as the metric and Ward’s criterion (83) as the linkage method. The optimal number of clusters was determined based on three criteria (84): the KL index, the Ball index, and the McClain and Rao index.

The first cluster grouped the frontoparietal, dorsal atteThe first cluster grouped the frontoparietal, dorsal attention, and somato/motor networks. Following recent recommendations on network correspondence (85), we computed similarities between the clusters’ topography and a set of task-defined networks (49, 50) using the Sørensen-Dice index(86, 87) (*SDI*). We then labeled this cluster as attention-related because these networks corresponded to the task-defined lateralized visuospatial attention network (*SDI*=0.33, Extended Data Fig. 5b), which was higher than their correspondence with the lateralized supramodal language network (49) (*SDI*=0.16, Extended Data Fig. 5a). Similarly, the second cluster was designated language-related as it showed a stronger correspondence with the task-defined language network (*SDI*=0.25) compared to the attention network (*SDI*=0.22).

We then examined the overall lateralization of biomarkers in the attention- and language-related clusters (Fig. 5d). Barnard’s unconditional test (88) revealed a significant difference in the distribution of leftward and rightward asymmetry indices between the two clusters (*p*=0.0047). The attention-related cluster exhibited a higher proportion of negative asymmetry indices (67%), indicating rightward asymmetry, whereas the language-related cluster showed a higher proportion of positive index (71%), indicating leftward asymmetry. An odds ratio of 5 indicated that networks related to attention were five times more likely to exhibit negative (rightward) asymmetry than those related to language. This significant association demonstrates that the two clusters have distinct asymmetrical patterns at the molecular and cellular levels. This result aligns with the literature demonstrating left-hemisphere anatomical and functional dominance for language and right-hemisphere dominance for visuospatial attention from birth (15). While the intrinsic functional networks were not the focus of our primary analyses, we used them as a canonical organizational scaffold to contextualize the spatial distribution of the molecular and cellular asymmetries identified. This provided an interpretable framework to summarize the embedding of lateralized biological features within established large-scale systems, without implying that resting-state networks directly drive, or are driven by, these asymmetries. Although speculative, these mitochondrial phenotypes and cellular correlates may serve as potential markers reflecting the functional lateralization of language and visuospatial attention. The distinct molecular and cellular asymmetries observed, with leftward bias in language-related networks and rightward bias in attention-related networks, reinforce the complementary nature of hemispheric specialization rooted in biological organization (36).

## Discussion

Through the integration of data across biophysical scales, including neurotransmitters, cell-types, mitochondrial phenotype, and functional neuroimaging data, we demonstrate that hemispheric specialization is a key principle of brain organization from the microstructural to the macroscale level, with two opposing lateralized systems supporting language and attentional processes. Our data revealed that functional lateralization is reflected in a neurochemical axis anchored by acetylcholine and norepinephrine (Fig. 2). While centered on cholinergic and noradrenergic asymmetries, this axis reveals a complex neurochemical architecture, with dopaminergic and serotonergic systems contributing meaningfully to its multivariate profile. A rightward lateralization of the muscarinic acetylcholine receptor (M_1_) associated to a leftward dominance of the norepinephrine receptor (NET) is linked to the dominance of the left hemisphere during language processing, while an opposite pattern was observed related to visuospatial attention processing. Of note, the multimodal monoaminergic-cholinergic was spatially coupled to the lateralization of both mitochondrial phenotypes (Fig. 3) and cellular subtypes (Fig. 4). Critically, the mitochondrial phenotypes and cellular subtypes formed a fingerprint discriminating between a left-lateralized system related to language processing and its right-lateralized counterpart supporting visuospatial attention processing (Fig. 5). Together, this work advances our understanding of the close relationship between the localized hemispheric specialization of specific behaviors and their underlying cellular and molecular substrates.

Our findings suggest a key role for neurotransmitter asymmetries, particularly along the multimodal monoaminergic-cholinergic axis, in shaping functional hemispheric specialization. Norepinephrine, predominantly produced by the locus coeruleus, has been extensively implicated in attentional control, working memory, and behavioral flexibility (89). It facilitates top-down regulation by the dorsolateral prefrontal cortex, supporting cognitive processes such as attentional shifting and executive control (90). Moreover, noradrenergic activation promotes neural resetting, dynamically reallocating neural resources and enabling perceptual reorientation (91–93), and it contributes to learning by adjusting neural dynamics across multiple timescales and organizational levels (94). Recent work in animal models further suggests that the locus coeruleus activity modulates cortical network topology, influencing the prioritization of associative versus sensory information processing (95). Such findings complement neuropsychological evidence that the right hemisphere plays a dominant role in spatial attentional reorienting and awareness (96), consistent with the right-lateralized noradrenergic system identified here. Similarly, acetylcholine is essential for prefrontal cortex function, with its depletion leading to severe cognitive impairments comparable to the loss of the cortex itself (97). Acetylcholine fine-tunes attentional selection and behavioral flexibility, reinforcing its complementary role alongside norepinephrine in guiding cognitive processing (98). Notably, receptor distribution analyses have revealed asymmetries in neurotransmitter densities across hemispheres, particularly in the inferior parietal lobule and language-related areas, further supporting the neurochemical underpinnings of functional specialization (34). These neurotransmitter-driven functional asymmetries may be intrinsically linked to developmental mechanisms underlying hemispheric differentiation. In support of this, a growing genetic literature shows that human hemispheric asymmetries relevant to language arise from early, genetically regulated left-right developmental programs, including microtubule-mediated mechanisms(99, 100). Loci associated with structural asymmetries in classical language regions, such as the planum temporale and inferior frontal cortex, suggest that individual variation in the neuromodulatory and mitochondrial asymmetries may partly reflect these human-specific genetic pathways (100). Acetylcholine’s role in neurite elongation (101) aligns with evidence of microtubule-mediated brain asymmetry, as suggested by genetic findings implicating the TUBB4B tubulin gene in handedness (102). Given the association between microtubule dynamics and atypical lateralization (103), one could predict that individuals with atypical hemispheric organization for language (4, 5, 104), might exhibit an inverted or more balanced pattern of neurotransmitter lateralization. Intriguingly, such neurochemical patterns are reminiscent of those observed in neuropsychiatric and neurodevelopmental disorders (105), illnesses marked by atypical lateralization patterns such as reduced cortical thickness and altered language network organisation (106) associated to polygenic risks in schizophrenia (107), or functional rightward lateralization of the language network in major depressive disorder (108, 109). This raises the question of why some individuals with atypical lateralization remain asymptomatic while others develop psychiatric symptoms, positioning right-lateralized language individuals as a valuable model for understanding the neurobiological foundations of psychiatric vulnerability. At a broader systems level, these asymmetries may contribute to shaping dynamic brain network organization, consistent with recent evidence that large-scale network dynamics reflect psychiatric illness status and transdiagnostic symptom profiles across health and disease (110). Finally, our results suggest that the somato/motor network plays a role in modulating the balance of the multimodal monoaminergic-cholinergic axis, consistent with the motor theory of the origin of language proposing that lateralized manual behaviors (*e*.*g*., handedness) shaped the neural architecture underlying language evolution (111).

Although the canonical neurotransmitter axis we identified is primarily shaped by the opposing asymmetries of M_1_ and NET receptors, dopaminergic markers (D_1_, DAT, and D_2_) also exhibited moderate positive loadings. This finding may appear counterintuitive given the opposing intracellular signaling cascades of D_1_ (Gs-coupled) and D_2_ (Gi-coupled) receptors (29). Their co-loading on the same canonical axis likely reflects shared spatial asymmetries rather than redundant functional roles. Indeed, the dopaminergic system has been implicated in lateralized cognitive functions such as language, attention, and motor control (112–114). A similar interpretive nuance applies to the identified asymmetry in NET, which is a presynaptic norepinephrine reuptake transporter rather than a postsynaptic receptor. While NET is involved in signal attenuation via synaptic clearance (115), its spatial distribution still reflects regional variation in noradrenergic tone and locus coeruleus projections. As such, NET PET imaging remains a valuable topographic marker of the noradrenergic system (29). Our analytical focus is on relative asymmetries in neuromodulatory marker distribution, not direct synaptic signaling, making NET a valid contributor to the observed axis. Nonetheless, future studies incorporating whole-brain PET maps of postsynaptic adrenergic receptors, such as α- and β-receptor subtypes (116), will be essential to more directly assess the specificity of the multimodal monoaminergic-cholinergic contrast and its role in shaping hemispheric specialization.

Variations in mitochondrial bioenergetics influence cognitive performance and may contribute to cognitive impairments in mitochondrial diseases, underscoring their role in both energy production and molecular signaling essential for healthy brain functions (45). Mitochondrial respiratory capacity aligns with evolutionary patterns, supporting the elevated energy demands of human-specific cognitive functions (117, 118). Notably, brain regions involved in executive functions exhibit high expression of neuromodulatory receptors and a strong dependence on glucose metabolism (118), with mitochondrial respiratory capacity emerging as a key mitochondrial phenotype linking the multimodal monoaminergic-cholinergic axis to functional hemispheric specialization. Beyond energy production, mitochondrial metabolism regulates neurodevelopment by influencing cortical expansion (119). Hemispheric specialization may optimize brain energy use (120), possibly emerging from functional hemispheric spatial competition (48), where visuospatial functions are lateralized to the right hemisphere as a consequence to the left lateralization of language processing (1). While fMRI studies support both independent and causal complementarity mechanisms (1), the lack of a handedness effect on spatial lateralization challenges a strictly causal model (121).

Glutamatergic excitatory neurons, especially those with intratelencephalic projections, exhibit substantial diversity in morphology, electrophysiology, and gene expression, contributing to cortical specialization (122). This diversity is pronounced in the supragranular layers of the human cortex, where transcriptomic subtypes show depth-dependent variation in dendritic architecture, firing properties, and connectivity patterns (122). These structural and functional gradients likely support areal and laminar specialization, providing a substrate for hemispheric differences in cognitive functions. Excitatory neurons establish functionally specific circuits through stereotyped connections with interneurons, thereby supporting regional specialization and functional differentiation (123). Language regions exhibit leftward asymmetries in superficial cortical layers (2 and 3), while visuospatial attention areas show rightward asymmetries in deeper layers (5 and 6), correlating with behavioral differences (72). Our findings align with this literature, highlighting asymmetries of intratelencephalic projecting neurons across layers 2, 3, 5, and 6, linked to the multimodal monoaminergic-cholinergic axis underlying functional lateralization. Additionally, distinct laminar gene expression profiles within the left hemisphere’s language network suggest molecular adaptations for specialized cognitive processing and highlight implications for conditions such as dyslexia and schizophrenia (44).

From an evolutionary perspective, functional asymmetries reflect a fundamental principle of human brain organization (8) and a conserved feature across metazoan nervous systems (9). In primates, cortical expansion has predominantly occurred within association territories rather than primary sensory areas, supporting cognitive functions that operate independently of direct sensory input (124, 125). The aggregation of functions into hemispheric asymmetries enhances neural efficiency, facilitating specialized and parallel neurocognitive processing (126–128). Structural adaptations in grey and white matter, particularly in the prefrontal cortex, have likely played a central role in human cognitive evolution, supporting complex functions such as language (129, 130). Beyond genetic influences, hemispheric asymmetries emerge from dynamic interactions between developmental, environmental, and pathological factors, giving rise to both typical and atypical phenotypes (36, 131). Population-level language lateralization is rooted in polygenic influences (132, 133), while interhemispheric connectivity, mediated by the corpus callosum, plays a critical role in functional specialization (4). This connectivity profoundly shapes cognitive domains such as language and executive function (10). At the network level, cognitive functions are supported by intracortical connectivity that follows a hierarchical organization, for instance as described by Fuster’s perception-action cycle model (134, 135). These principles underlie large-scale cognitive networks (33), including those implicated in language (49, 136) and attention (50).

Several potential limitations should be considered when interpreting the present findings. First, while our study elucidates potential cellular and molecular substrates underlying hemispheric specialization, the causal pathways linking lateralized cognitive functions to these biological mechanisms remain to be established. Additionally, the cross-sectional nature of the data precludes inferences about the developmental trajectory through which hemispheric specialization, cellular and molecular substrates, and global brain architecture interact over time. Thus, the dynamic interplay between lateralized functions and brain-wide organization remains an open question. Methodologically, we accounted for potential variance introduced by left-handed and atypically lateralized individuals in our task-based activation analyses; however, the presence of such individuals in our cellular, molecular, and neurotransmitter datasets could not be determined and may contribute to variability. Nevertheless, receptor fingerprints remain relatively stable across individuals, suggesting that group-level analyses may still provide meaningful insights (31). However, future studies should prioritize individual-level approaches, particularly in the context of personalized medicine (137, 138). At the opposite extreme, our mitochondrial phenotype data lack variability, being derived from a single neurotypical brain sample (46), underscoring the need for larger, more diverse datasets. Furthermore, although the use of PCA followed by CCA was methodologically motivated to reduce collinearity and enhance interpretability, we recognize that this dimensionality reduction constrains the analysis to operate along the dominant axes of variation in the data. While this approach is appropriate given the high autocorrelation between language modalities and across neurotransmitter maps, it reinforces the exploratory nature of our findings. Nevertheless, the observed associations provide specific and testable hypotheses for future confirmatory studies in independent and individual-level datasets. Furthermore, our analyses focus on individuals with presumed typical hemispheric specialization, who represent approximately 92% of the population (5). However, we acknowledge the existence of individuals with atypical language lateralization (104), whose brain organization does not simply mirror that of typically lateralized individuals (4). Investigating this specific subgroup, including cases of co-dominant language function (139), is essential for a more comprehensive understanding of how cellular and molecular mechanisms contribute to hemispheric specialization. This issue resonates with a broader call in psychiatry to embrace variability as an essential feature rather than a confound (140). Finally, our study is constrained by the spatial limitations of our datasets: imputed cell-type abundances from bulk-tissue microarray data provide limited spatial resolution, potentially underestimating true cell type-function relationships (41). The same limitation applies to neurotransmitter PET data, which would benefit from higher-resolution imaging in future research (29). Lastly, our analyses were restricted to the cortex, omitting potential subcortical contributions to hemispheric specialization. Nevertheless, these results provide a foundation for future brain-constrained models that integrate hemispheric specialization as a fundamental property of brain organization (138).

The present findings demonstrate that the hemispheric complementarity of language and visuospatial attention is mediated by the interhemispheric balance of an multimodal monoaminergic-cholinergic axis. Moreover, this axis is coupled with asymmetries of mitochondrial phenotypes and cell-type densities. Notably, the molecular and cellular densities collectively form a distinct fingerprint, differentiating a left-lateralized system associated with language processing from its right-lateralized counterpart supporting visuospatial attention. Together, these discoveries establish a direct connection between functional cortical specialization and its underlying neurotransmitter, molecular, and cellular architecture, offering critical insights into the mechanisms of cerebral dominance and their implications for cognition in health and disease.

## Methods

### Brain Maps

#### Functional Homotopic Atlas

To characterize the multimodal relationships among functional data, neurotransmitter density maps, and molecular and cellular biomarkers of the cortical sheet, we conducted regional analyses using the Atlas of Intrinsic Connectivity of Homotopic Areas (AICHA (52)). AICHA is a homotopic atlas (33, 141) optimized for analyzing functional data and specifically designed to assess hemispheric lateralization. It comprises 163 cortical homotopic brain regions in each hemisphere.

The hemispheric differences, *i*.*e*. asymmetry maps, for each modality were defined as *left* - *right*: we subtracted the regional values of the right hemisphere from those of the left hemisphere, and thus, a positive value means a leftward asymmetry (*left* > *right*), resulting in lateralization maps across 163 regions. As highlighted by the Laterality Indices Consensus Initiative (142), no universal consensus exists on the optimal laterality index formula; authors are therefore encouraged to clearly define their metric, which we do here. The *left* - *right* difference has been recommended for its robustness to noise (142) and preprocessing effects (143, 144) and for preserving full information without thresholding (142). In contrast, normalized indices have the potential to amplify differences or introduce denominator discontinuities (145) and yield results highly similar to the non-normalized approach (2). Moreover, direct *left* - *right* statistical comparison methods have demonstrated superior correspondence with the gold-standard Wada test for determining hemispheric language dominance (146). Additionally, the *left* - *right* method avoids misclassifying bilateral patterns (147), while ensuring methodological consistency with a growing body of recent work employing similar definitions of hemispheric asymmetry (4, 5, 49, 50, 58, 128, 136, 148, 149).

#### fMRI Activation Maps

Functional brain lateralization was assessed using four previously published maps encompassing the two major and most widely described lateralized cognitive functions: language and visuospatial attention. These maps represent average cortical activation in 125 strongly typical participants for language (38% female, *μ*_Age_=25.73 years, 46% left-handers), covering activation in 326 cortical brain regions from the AICHA atlas. Seven region pairs in the orbital and inferior temporal regions were excluded due to signal reduction from susceptibility artifacts. Then, we computed asymmetry maps, resulting in 163 asymmetric regional scores (Fig. 1a). Functional maps are provided in MNI stereotaxic space (MNI ICBM 152, template sampling size of 2×2×2 mm^3^ voxels; bounding box, *x*=−90 to 90 mm, *y*=−126 to 91 mm, *z*=−72 to 109 mm). We focused on strongly typical individuals, who reliably (4, 5) exhibit the canonical pattern of hemispheric specialization: with dominant left-hemisphere engagement for language (49) and right-hemisphere dominance for visuospatial attention (50). This group shows minimal variability in hemispheric complementarity, thereby providing a stable model for investigating the molecular and cellular correlates of functional lateralization. The four maps included three language-related contrasts (sentence production minus an overlearned word list, listening, and reading) and a visuospatial attention contrast using a line bisection judgment task.

The protocols, structural and functional image acquisition parameters, and image analysis procedures for each task have been previously detailed (49, 50). All four tasks are part of the BIL&GIN database (51). Images were acquired using a 3T Philips Intera Achieva scanner (Philips, Eindhoven, The Netherlands). Task-related functional volumes were obtained using a T2*-weighted echo-planar imaging sequence (T2*-EPI; TR=2 s; TE=35 ms; flip angle=80°; 31 axial slices; field of view=240 × 240 mm^2^; voxel size=3.75×3.75×3.75 mm^3^ isotropic; 240 volumes). The first four volumes of each sequence were discarded to allow for MR signal stabilization. Functional data were processed using global linear modeling with Statistical Parametric Mapping software (SPM12, www.fil.ion.ucl.ac.uk/spm/). The three language-related maps consisted of three runs comparing sentence tasks (involving phonological, semantic, prosodic, and syntactic processing) with a word-list reference task, which is less complex but still high-level verbal processing. Each run included 13 blocks of the sentence task and 13 blocks of the word-list task. The reading and listening runs were 6 minutes and 28 seconds long, while the production task was 6 minutes and 24 seconds. The attention task comprised 36 trials alternating the presentation of a horizontal line bisected by a short vertical line biased toward the left, right, or centered, followed by a fixation cross; this run lasted 8 minutes.

#### Neurotransmitter Systems Maps

Receptor density data were obtained from a report (29) having collected data from multiple PET studies involving 1,238 healthy adults (42% female, *μ*_weigted Age_=36.56 years). This report provides the density of 19 neurotransmitter receptors and transporters, which were parcellated into 326 cortical regions of AICHA, the same atlas used to parcel functional MRI data. These regional values were standardized within each map before computing asymmetries (Fig. 1b). Before regional analysis, neurotransmitter maps were linearly registered to the MNI stereotaxic space (MNI ICBM 152) using the R library *RNiftyReg* (150) (R package version: 2.8.1). Following Hansen and colleagues’ procedure (29), we combined using weighted average neurotransmitter maps with more than one mean image of the same tracer: 5-HT_1B_, D_2_, mGluR_5_ and VAChT. Altogether, the images are an estimate proportional to receptor densities, and we, therefore, refer to the measured value (that is, binding potential and tracer distribution volume) simply as density.

#### Synaptic Density Map

Receptor density data were obtained from Johansen and colleagues (40), which collected data using [^11^C]UCB-J PET imaging that binds to Synaptic Vesicle Glycoprotein 2A (SV2A), a presynaptic protein considered a reliable proxy for synaptic density. The study involved 33 healthy participants (17 females; *μ*_median Age_=24 years), and regional binding values were calibrated to absolute SV2A densities (pmol/mL) using postmortem human autoradiography data. PET scans were processed with a standardized surface- and volume-based pipeline, aligning images to MNI ICBM 152 volume space. We then parcellated the SV2A map into 326 cortical regions of AICHA, the same atlas used to parcel functional MRI and neurotransmitter density data. These regional values were standardized before computing asymmetries (Fig. 3a).

#### Mitochondrial Phenotypes Maps

Mitochondrial phenotype data were obtained from Mosharov and colleagues (46). The frozen right hemisphere of a 54-year-old neurotypical healthy male was physically voxelized into 703 cubes, each measuring 3×3×3 mm, yielding a resolution comparable to standard neuroimaging. Markers of mitochondrial content (CS^1/3^ and mtDNA^1/3^) were combined into a single metric, termed MitoD, to represent the mitochondrial tissue density within each voxel. Enzymatic activities of oxidative phosphorylation (OxPhos) were measured via colorimetric and respirometry assays and averaged to generate a tissue respiratory capacity map (TRC), reflecting the overall OxPhos capacity per milligram of tissue. Finally, TRC was expressed relative to MitoD (MRC=TRC/MitoD) to produce the mitochondrial respiratory capacity map, indicating tissue-specific mitochondrial specialization for oxidative metabolism on a voxel-by-voxel basis.

These three maps were then linearly registered to the MNI stereotaxic space (MNI ICBM 152) using the R library *RNiftyReg* (150) (R package version: 2.8.1). We parcellated the three maps into 326 cortical regions of AICHA, the same atlas used to parcel functional MRI, neurotransmitter density and synaptic density data. These regional values were standardized before computing asymmetries (Fig. 3a).

#### Cell-Type Abundance Maps

Cell-type maps were obtained from Zhang and colleagues (41). Cell-type maps were derived using the Allen Human Brain Atlas transcriptional data and single-nucleus RNA-sequencing data from eight cortical regions. Postmortem bulk-tissue gene expression data from the Allen Human Brain Atlas were deconvolved using a validated method to estimate cell-type abundances. This approach leveraged transcriptional signatures from single-nucleus RNA-sequencing datasets to identify 24 distinct cellular classes (151). These included nine GABAergic interneurons, nine glutamatergic excitatory neurons, and six non-neuronal cell-types. The cell-type abundances for each Allen Human Brain Atlas cortical sample were mapped to the cortical voxel represented in MNI ICBM 152 volume space and then parceled into the AICHA atlas. Samples were first averaged within parcels at the individual donor level and then averaged across donors. These regional values were standardized before computing asymmetries. A visualization of the 24 cell-type asymmetry maps is available in Extended Data Fig. 4. Due to the sampling procedure, the cell-type asymmetry maps only included 123 pairs of homotopic cortical regions.

### Statistical Analyses

Statistical analyses were performed using R (152) (R version: 4.3.3). The neuroimaging data were handled and processed using R libraries *neurobase* (153) (R package version: 1.32.4), *RNifti* (154) (R package version: 1.7.0), and *RNiftyReg* (150) (R package version: 2.8.1). Data wrangling was performed using the R library *dplyr* (155) (R package version: 1.1.4). Brain visualizations were realized using Surf Ice (156, 157) (version: 6 october 2021), and were made reproducible following guidelines to generate programmatic neuroimaging visualizations (158).

#### Canonical Correlation Analysis to Assess the Functional Cortical Lateralization – Neurotransmitters Density Asymmetries Associations

In order to clarify the potential links between cortical lateralization and asymmetries in neurotransmitter density, we quantified the linear associations using Canonical Correlation Analysis (CCA) approach (53) (R library *acca* (159), version: 0.2). As a multivariate technique, CCA identifies optimal linear combinations from two distinct sets of variables to maximize their mutual correlation, thereby revealing modes of joint variation that highlight the interplay between hemispheric dominance and the lateralization of its neurotransmitter correlates. The analysis yielded a series of m orthogonal canonical modes, with each mode capturing a distinct portion of the shared variance between functional lateralization and neurotransmitter density that is not explained by the remaining modes.

Prior to the CCA, we performed a dimensionality reduction on our high-dimensional brain dataset, i.e. comprising functional tasks and neurotransmitter density maps asymmetries, via principal component analysis (53) (PCA). We retained components selected based on the elbow criterion, which reflects the variance explained by successive components, using the *PCAtools* R package (160) (version 2.5.15). The resulting principal components served respectively as cortical lateralization set and neurotransmitter asymmetries set for the subsequent CCA.

To assess mode significance, we used a spin test. In this procedure, the cortical regions as defined in the AICHA atlas were rotated along an inflated surface to generate 10,000 null atlas configurations that preserve the cortex’s intrinsic spatial autocorrelation (56). These null configurations were then used to shuffle the rows of our feature data frame, i.e. the asymmetry maps of the four tasks, thereby creating a null distribution of mode correlation values. The statistical significance (denoted as *p*_spin_) was determined as the proportion of null values exceeding the observed correlation. Null maps were generated using the *netneurotools* Python toolbox (release 0.2.5, Python 3.12.5; github.com/netneurolab/netneurotools). To further identify which functional and neurotransmitter maps contributed most to the task-fMRI and PET asymmetry modes, we quantified canonical loadings as product-moment correlations between each input map and its corresponding canonical variate. Their significance was evaluated via bootstrapping (10,000 resamples), using the ratio of the observed correlation to its bootstrap-derived standard error to compute *z*-scores and corresponding two-tailed *p*-values (161). Loadings were considered reliable after Holm family-wise error correction across input features (*p*_FWER_<0.05).

#### Identification of Molecular Biomarkers Spatially Correlated to the Multimodal Monoaminergic-Cholinergic Axis

To identify potential relationships between the neurotransmitter multimodal monoaminergic-cholinergic axis and molecular biomarkers (three mitochondrial phenotypes maps (46): tissue respiratory capacity, mitochondrial tissue density, and mitochondrial respiratory capacity, and a synaptic activity map: synaptic vesicle glycoprotein 2A (40)) we used linear regression. The multimodal monoaminergic-cholinergic axis is the dependent variable. To assess significance of each relationship, we used a spin test (10,000 permutations). *P*-values were corrected for multiple comparisons using the false discovery rate (FDR) method (68) (denoted as *p*_FDR_), accounting for the four regression models applied separately for each molecular map modeling the association with the multimodal monoaminergic-cholinergic axis.

#### Multimodal Cellular Underpinning of the Multimodal Monoaminergic-Cholinergic Axis

Due to the multidimensional nature of the relationship between large-scale functional organization and cellular subtype expression (41), we performed stepwise multiple linear regression to assess the association between the multimodal monoaminergic-cholinergic axis and the asymmetry in the density of 24 cellular subtypes (41) (Extended Data Fig. 4). Stepwise regression was used to select the cellular subtypes that minimized the Akaike Information Criterion, yielding the most parsimonious model while maximizing the model likelihood associated with the multimodal monoaminergic-cholinergic axis. To assess significance of each selected cellular subtype, we used a spin test (10,000 permutations). *P*-values were corrected for multiple comparisons using FDR.

#### Mitochondrial and Cellular Fingerprints of Large-Scale Networks

We characterized each large-scale intrinsic network, as defined by Yan and colleagues (81), according to its molecular asymmetry values (TRC, MitoD, and MRC) and cellular asymmetry values (L5 IT, Micro/ PVM, L6 IT, and L2/3 IT) that are significantly associated with the neurotransmitter asymmetry axis underlying functional lateralization. This approach yielded a unique fingerprint for each cerebral network (34). Each network was characterized by its normalized values for the seven biomarkers.

Based on these fingerprints, we tested the following hypothesis: do large-scale intrinsic cerebral networks follow a general principle of lateralization? Following the method defined by Zilles and colleagues (34), we investigated the presence of clusters among the seven networks using agglomerative hierarchical clustering (82), employing Euclidean distance as the metric and Ward’s criterion (83) as the linkage method. The optimal number of clusters was determined based on three criteria (84) using the *NbClust* package (R package version: 3.0.1): the KL index, the Ball index, and the McClain and Rao index. Subsequently, we labelled the identified clusters following recent recommendations on network correspondence (85). We computed similarities between the clusters’ topography and a set of task-defined networks (49, 50) using the Sørensen-Dice index (86, 87) (SDI).

Finally, we examined the lateralization of biomarkers within the identified clusters. Using Barnard’s unconditional test (88) (implemented via the *Barnard* R package (162), version: 1.8), we tested the significance of differences in lateralization between the clusters.

## Data and Code Availability Statement

The data (including: the average brain activity maps during three language tasks and one visuospatial attention task, the 19 neurotransmitter systems maps, the 24 cell-type maps, the synaptic density map, and the three mitochondrial phenotypes maps) and the code used in the Method section to process the data, to reproduce the results and visualizations can be found here (163): github.com/loicla-bache/Labache_2025_MolCelLat.

## Acknowledgments

This work was supported by the National Institute of Mental Health (R01MH120080 to AJH).

## CRediT Authorship Contribution Statement

**Loïc Labache:** Conceptualization, Data Curation, Formal Analysis, Investigation, Methodology, Resources, Software, Validation, Visualization, Writing—original draft, Writing—review & editing, Supervision, and Project administration. **Sidhant Chopra:** Methodology, Validation, Writing—review & editing. **Xi-Han Zhang:** Resources, Validation, Writing—review & editing. **Avram J. Holmes:** Conceptualization, Funding acquisition, Investigation, Methodology, Project administration, Resources, Supervision, Validation, Visualization, Writing—original draft, Writing—review & editing.

## Extended Data Figures

**Extended Data Fig. 1.**
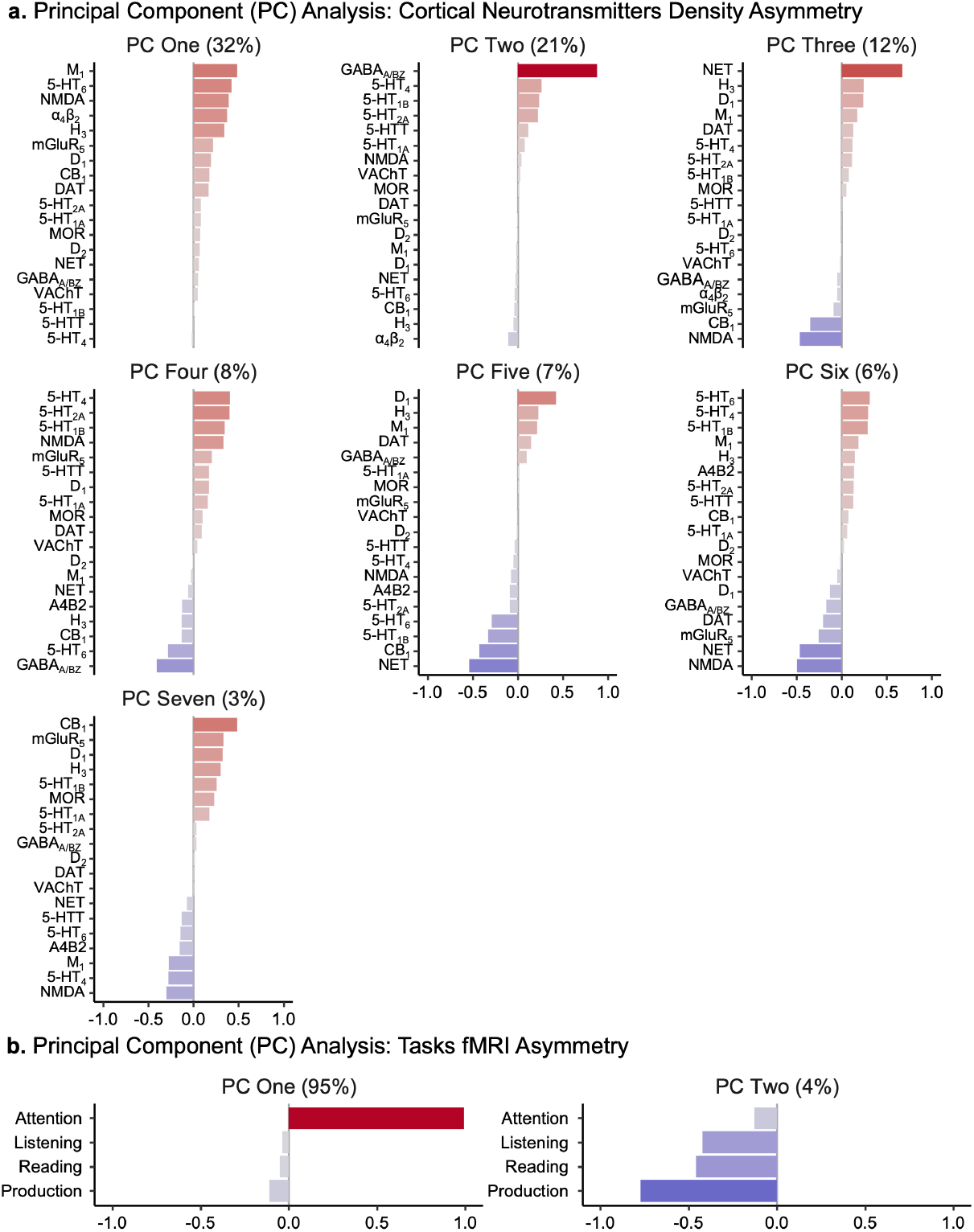
Principal Component Analysis (PCA) to produce a parsimonious and interpretable description of the original datasets (neurotransmitter asymmetries and cortical lateralization sets). **a**, Results of the PCA performed on the 19 cortical neurotransmitters density asymmetry maps (neurotransmitter asymmetries set). Based on the elbow criterion, we retained seven components, explaining 89% of the original data variances. The first three components explained 62% of the total variance, and can be decomposed as follow according to their loadings: PC One (32%) indicated how leftward lateralized acetylcholine is, PC Two (21%) indicated how much left asymmetrical GABA neurotransmitter are, and PC Three (12%) is related to the balance between norepinephrine and glutamate neurotransmitters. **b**, Results of the PCA performed on the 4 functional asymmetry tasks maps (cortical lateralization set). Based on the elbow criterion, we retained two components, explaining 99% of the original data variances. These two components can be decomposed as follow according to their loadings: positive loadings for PC One (95%) indicated strong leftward asymmetries displayed during the attention task, while negative loadings are linked to rightward asymmetries. Positive loadings for PC Two (4%) indicated strong rightward asymmetries displayed during the thress language related tasks, while negative loadings are linked to leftward asymmetries.

**Extended Data Fig. 2.**
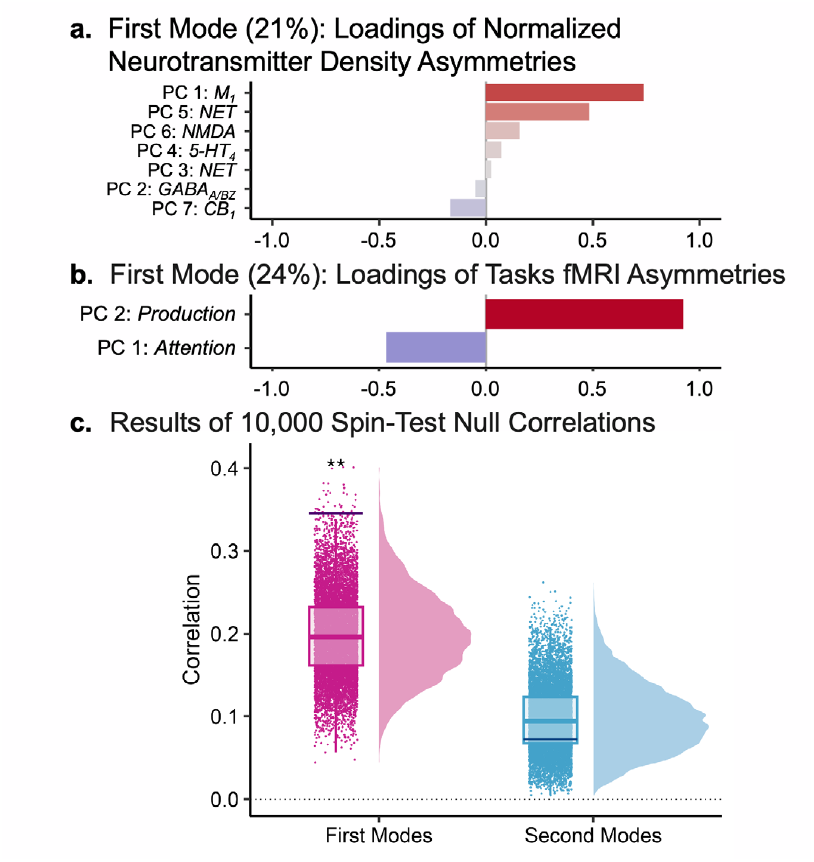
Canonical Correlation Analysis (CCA) results. **a**, Loadings of normalized neurotransmitter density asymmetries for the first CCA mode. This mode explained 21% of the variance and was led by the first principal component, indicating the degree to which acetylcholine neurotransmitters (M_1_) are leftward lateralized. **b**, Loadings of task fMRI asymmetries for the first CCA mode. This mode explained 24% of the variance and indicated the balance between the second and first components of the cortical lateralization set. This mode reflects hemispheric complementarity between language and attention tasks. **c**, Raincloud plots displaying results of the spin tests assessing the significance of the correlation between the neurotransmitter mode and the task mode. The spin test was performed using 10,000 permutations.

**Extended Data Fig. 3.**
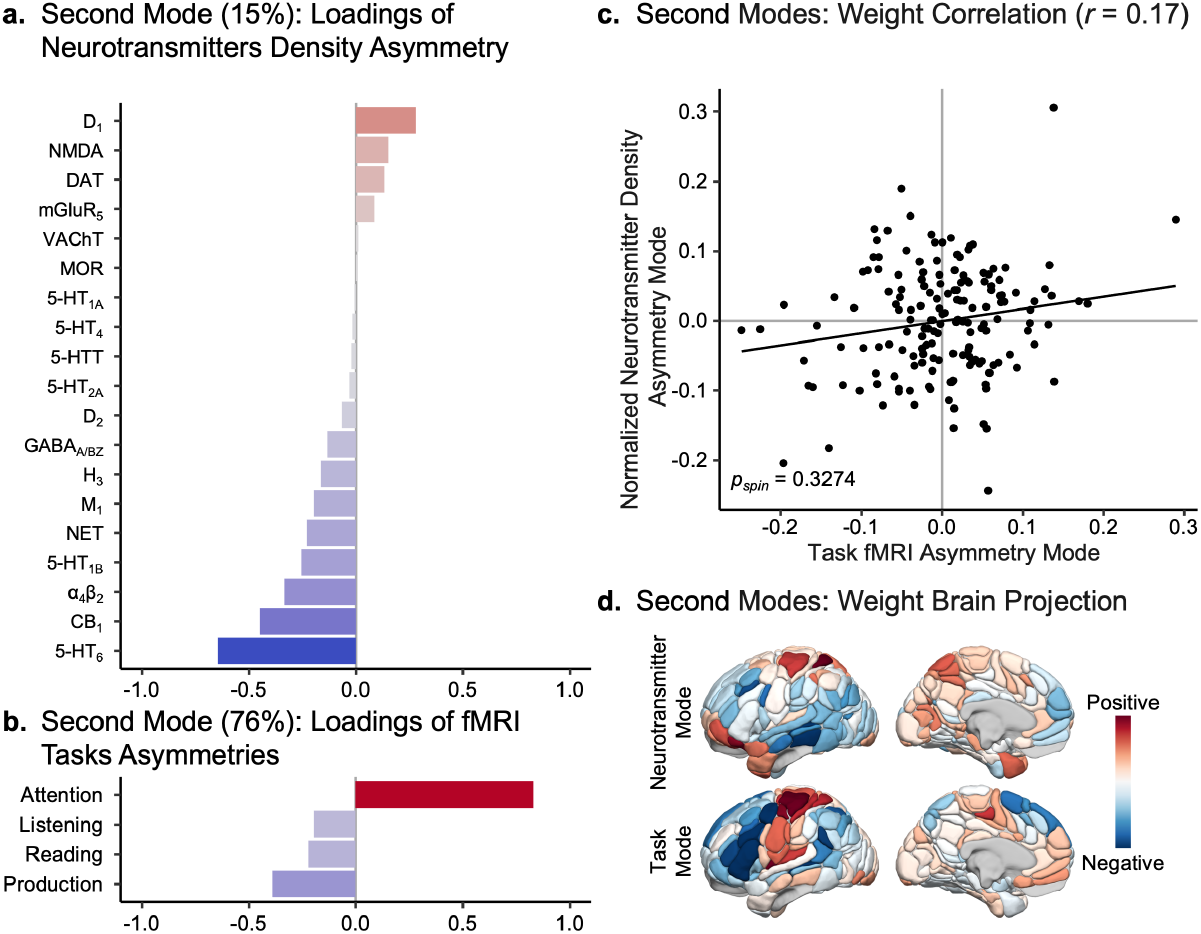
Second mode of the Canonical Correlation Analysis (CCA). **a**, Loadings values of the second neurotransmitter mode. This neurotransmitter mode explained 15% of the variance in the normalized neurotransmitter density asymmetry maps. **b**, Loadings values of the second task mode. **c**, Scatter plot showing the association between the second neurotransmitter and second task modes. The second neurotransmitter and task modes are not significantly correlated (*r*=0.17, *p*_*spin*_=0.3274; significance was assessed using a spin test with 10,000 permutations^52^). **d**, Projection of each regional CCA weight. The 3D rendering in the MNI space was created using Surf Ice software (https://www.nitrc.org/projects/surfice/).

**Extended Data Fig. 4.**
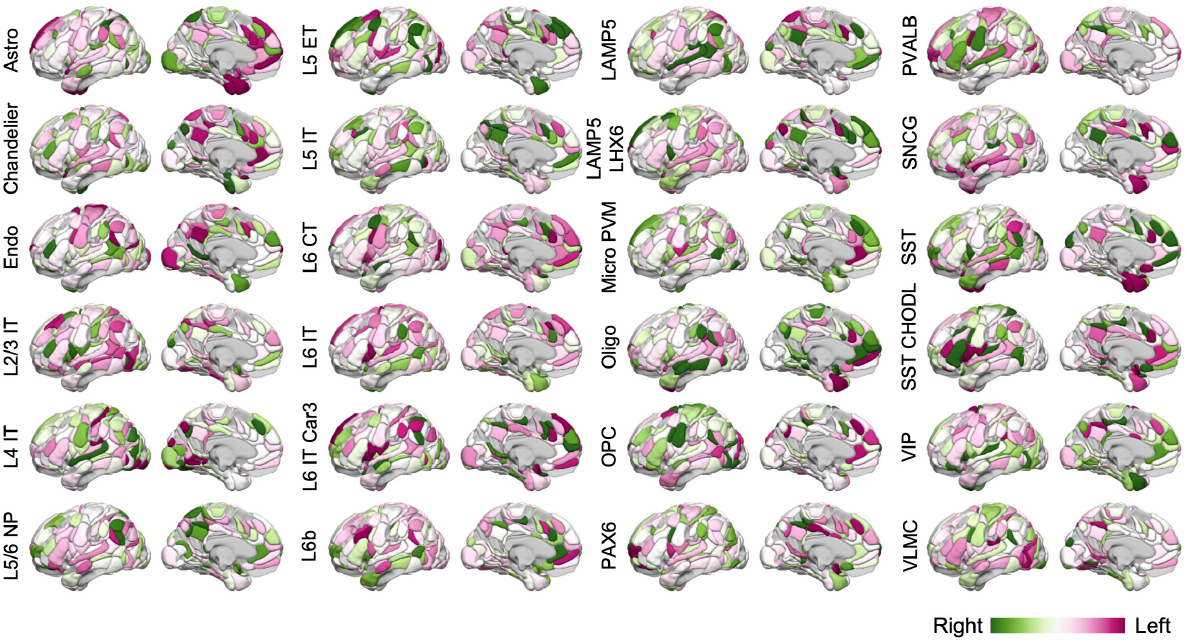
Normalized asymmetry maps of cellular biomarkers. Set of 24 normalized asymmetry maps of cellular biomarkers as reported by Zhang and colleagues^33^, derived from bulk samples of six postmortem brains^71^. Asymmetries were calculated by subtracting left hemisphere values from right hemisphere values (Right - Left) and are visualized on the left hemisphere. The maps encompass 123 cortical regions defined by the homotopic AICHA atlas^44^.

**Extended Data Fig. 5.**
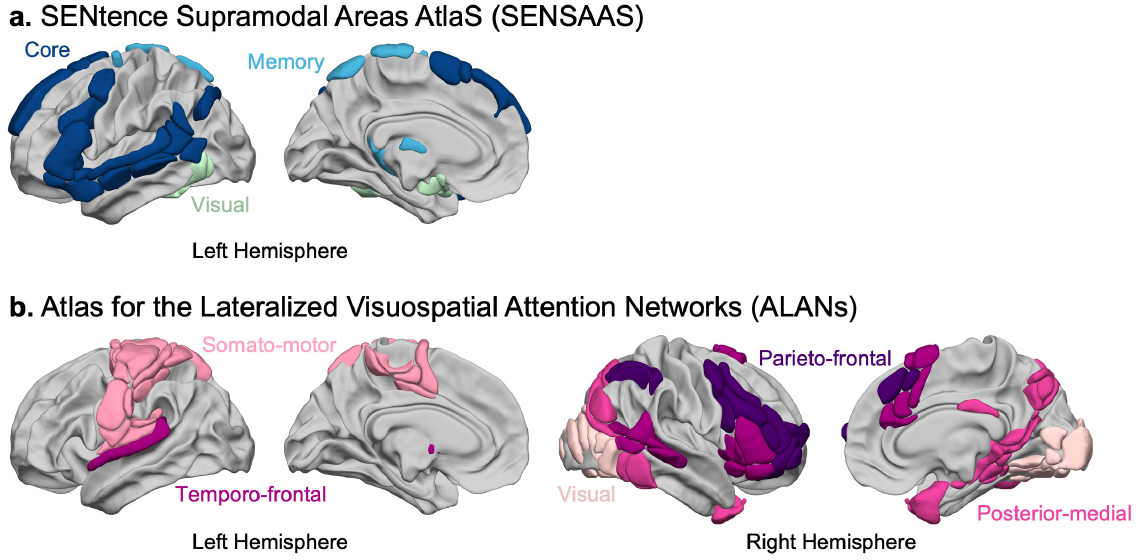
Functional atlases of lateralized cognitive functions. **a**, The sentence supramodal areas atlas^41^ comprises three networks that provide the anatomo-functional support for sentence processing. The dark blue network (Core) corresponds to the essential core language network, where lesions in its regions lead to aphasia. The blue network (Memory) represents a language-support network involved in episodic memory. The light blue network (Visual) represents a language-support network involved in visual processing. **b**, The atlas for the lateralized visuospatial attention networks comprises five networks that provide the anatomo-functional support for visual-spatial attention processing^42^. This atlas includes two primary core attentional networks: the parieto-frontal (dark purple) and temporo-frontal (purple) networks, as well as three support networks: the posterior-medial (light pink) network, and the visual and somato-motor networks, both of which support sensory-motor functions during visuospatial attention tasks.

## Extended Data Tables

**Extended Data Table 1.**
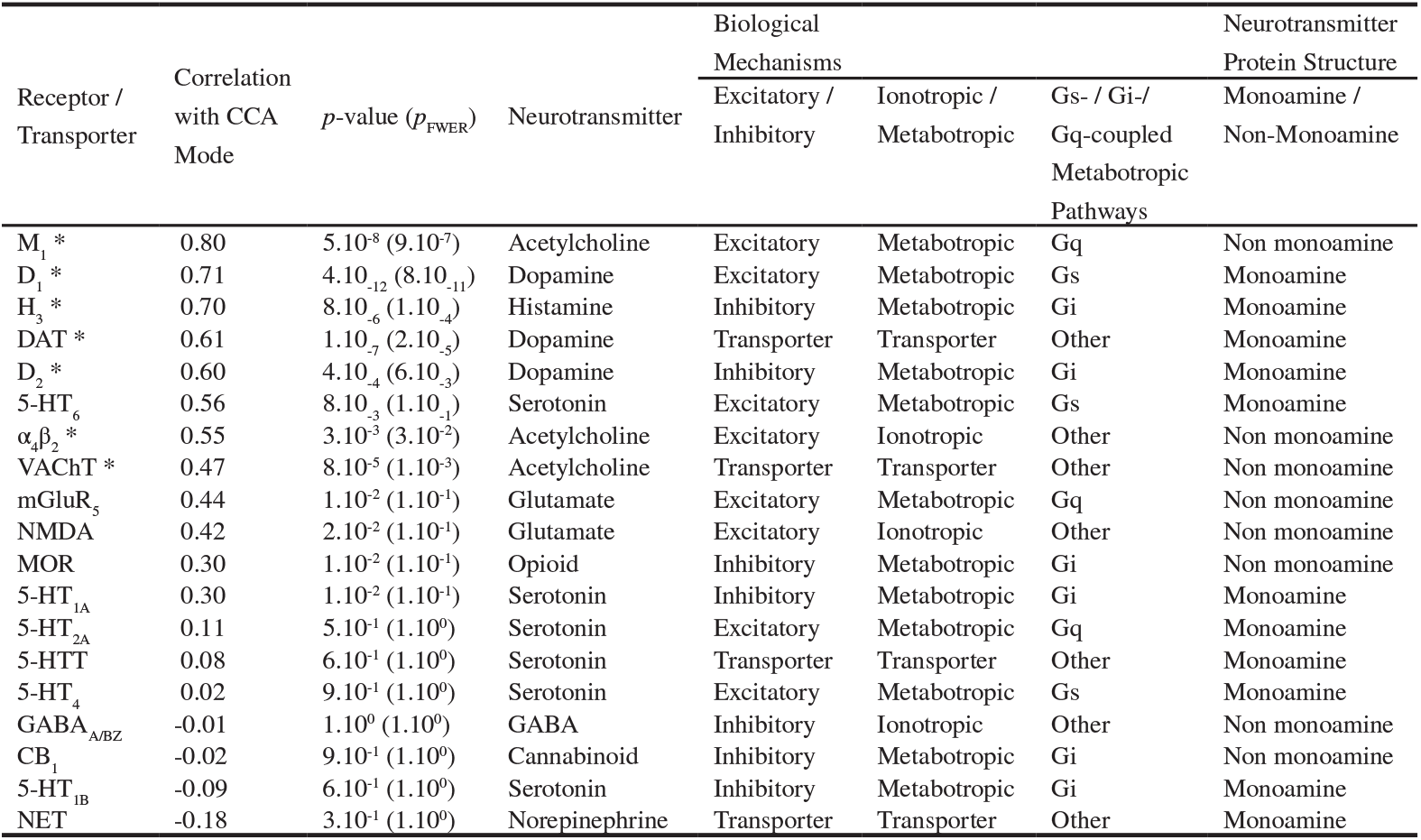
Correlation values and associated *p*-values of the neurotransmitter mode with the 19 normalized neurotransmitter density asymmetries. Neurotransmitters are stratified by biological mechanism (excitatory/inhibitory; ionotropic/metabotropic; Gs/Gi/Gq-coupled) and neurotransmitter class (monoamine/non-monoamine) as described by Hansen and colleagues (29). Asterisks (*) in the Receptor/Transporter column indicated statistical significance using bootstrapping (*p*_FWER_<5.10^−2^).

| Receptor /<br>Transporter | Correlation<br>with CCA<br>Mode | $p$-value ($p_{\text{FWER}}$) | Neurotransmitter | Biological<br>Mechanisms | | | Neurotransmitter<br>Protein Structure |
| --- | --- | --- | --- | --- | --- | --- | --- |
|  |  |  |  | Excitatory /<br>Inhibitory | Ionotropic /<br>Metabotropic | Gs- / Gi-/<br>Gq-coupled<br>Metabotropic<br>Pathways | Monoamine /<br>Non-Monoamine |
| M<sub>1</sub> * | 0.80 | 5.10<sup>-8</sup> (9.10<sup>-7</sup>) | Acetylcholine | Excitatory | Metabotropic | Gq | Non monoamine |
| D<sub>1</sub> * | 0.71 | 4.10<sup>-12</sup> (8.10<sup>-11</sup>) | Dopamine | Excitatory | Metabotropic | Gs | Monoamine |
| H<sub>3</sub> * | 0.70 | 8.10<sup>-6</sup> (1.10<sup>-4</sup>) | Histamine | Inhibitory | Metabotropic | Gi | Monoamine |
| DAT * | 0.61 | 1.10<sup>-7</sup> (2.10<sup>-5</sup>) | Dopamine | Transporter | Transporter | Other | Monoamine |
| D<sub>2</sub> * | 0.60 | 4.10<sup>-4</sup> (6.10<sup>-3</sup>) | Dopamine | Inhibitory | Metabotropic | Gi | Monoamine |
| 5-HT<sub>6</sub> | 0.56 | 8.10<sup>-3</sup> (1.10<sup>-1</sup>) | Serotonin | Excitatory | Metabotropic | Gs | Monoamine |
| α<sub>4</sub>β<sub>2</sub> * | 0.55 | 3.10<sup>-3</sup> (3.10<sup>-2</sup>) | Acetylcholine | Excitatory | Ionotropic | Other | Non monoamine |
| VACHT * | 0.47 | 8.10<sup>-5</sup> (1.10<sup>-3</sup>) | Acetylcholine | Transporter | Transporter | Other | Non monoamine |
| mGluR<sub>5</sub> | 0.44 | 1.10<sup>-2</sup> (1.10<sup>-1</sup>) | Glutamate | Excitatory | Metabotropic | Gq | Non monoamine |
| NMDA | 0.42 | 2.10<sup>-2</sup> (1.10<sup>-1</sup>) | Glutamate | Excitatory | Ionotropic | Other | Non monoamine |
| MOR | 0.30 | 1.10<sup>-2</sup> (1.10<sup>-1</sup>) | Opioid | Inhibitory | Metabotropic | Gi | Non monoamine |
| 5-HT<sub>1A</sub> | 0.30 | 1.10<sup>-2</sup> (1.10<sup>-1</sup>) | Serotonin | Inhibitory | Metabotropic | Gi | Monoamine |
| 5-HT<sub>2A</sub> | 0.11 | 5.10<sup>-1</sup> (1.10<sup>0</sup>) | Serotonin | Excitatory | Metabotropic | Gq | Monoamine |
| 5-HTT | 0.08 | 6.10<sup>-1</sup> (1.10<sup>0</sup>) | Serotonin | Transporter | Transporter | Other | Monoamine |
| 5-HT<sub>4</sub> | 0.02 | 9.10<sup>-1</sup> (1.10<sup>0</sup>) | Serotonin | Excitatory | Metabotropic | Gs | Monoamine |
| GABA<sub>A/BZ</sub> | -0.01 | 1.10<sup>0</sup> (1.10<sup>0</sup>) | GABA | Inhibitory | Ionotropic | Other | Non monoamine |
| CB<sub>1</sub> | -0.02 | 9.10<sup>-1</sup> (1.10<sup>0</sup>) | Cannabinoid | Inhibitory | Metabotropic | Gi | Non monoamine |
| 5-HT<sub>1B</sub> | -0.09 | 6.10<sup>-1</sup> (1.10<sup>0</sup>) | Serotonin | Inhibitory | Metabotropic | Gi | Monoamine |
| NET | -0.18 | 3.10<sup>-1</sup> (1.10<sup>0</sup>) | Norepinephrine | Transporter | Transporter | Other | Monoamine |

**Extended Data Table 2.**
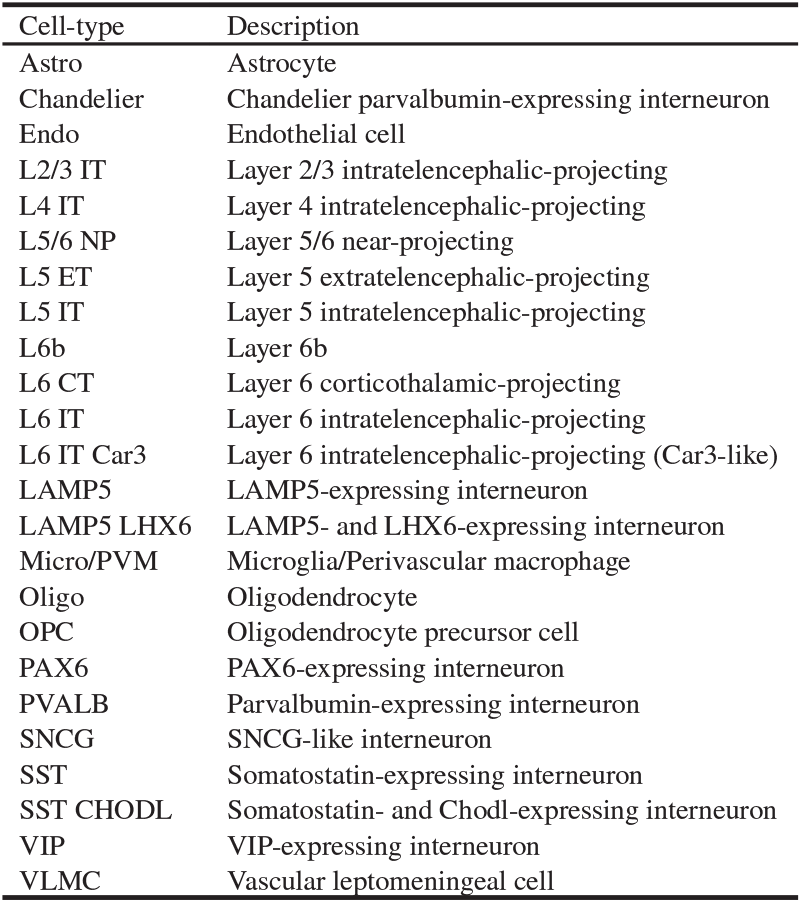
Cell-type annotations (41).

